# Origin and persistence of polymorphism in loci targeted by disassortative preference: a general model

**DOI:** 10.1101/2022.03.09.483509

**Authors:** Camille Coron, Manon Costa, Hélène Leman, Violaine Llaurens, Charline Smadi

**Affiliations:** Université Paris-Saclay, CNRS, Laboratoire de mathématiques d’Orsay, 91405, Orsay, France; Institut de Mathématiques de Toulouse; UMR5219. Université de Toulouse; CNRS. UPS, F-31062 Toulouse Cedex 9, France; Univ. Lyon, Inria, ENSL, UMPA, CNRS UMR 5669, 69364 Lyon, France; Institut de Systématique, Evolution et Biodiversité (UMR 7205 CNRS/MNHN/SU/EPHE/UA), Muséum National d’Histoire Naturelle - CP50, 57 rue Cuvier, 75005 PARIS, FRANCE; Univ. Grenoble Alpes, INRAE, LESSEM, 38000 Grenoble, France and Univ. Grenoble Alpes, CNRS, Institut Fourier, 38000 Grenoble, France

**Keywords:** heterogamy, overdominance, allelic differentiation, allelic turn-over

## Abstract

The emergence and persistence of polymorphism within populations generally requires specific selective regimes. Here, we develop an unifying theoretical framework to explore how disassortative mating can generate and maintain polymorphism at the targeted loci. To this aim, we model the dynamics of alleles at a single locus *A* in a population of haploid individuals, where reproductive success depends on the combination of alleles carried by the parents at locus *A*. Our theoretical study of the model confirms that the conditions for the persistence of a given level of allelic polymorphism depend on the relative reproductive advantages among pairs of individuals. Interestingly, equilibria with unbalanced allelic frequencies were shown to emerge from successive introduction of mutants.We then investigate the role of the function linking allelic divergence to reproductive advantage on the evolutionary fate of alleles within population. Our results highlight the significance of the shape of this function on both the number of alleles maintained and their level of genetic divergence. Large number of alleles are maintained with substantial turn-over among alleles when disassortative advantage slowly increases when allelic differentiation becomes large. In contrast, few highly differentiated alleles are predicted to be maintained when genetic differentiation has a strong effect on disassortative advantage. These opposite effects predicted by our model shed light on the levels of allelic differentiation and polymorphism empirically observed in loci targeted by disassortative mate choice.

## 1 Introduction

Selective mechanisms favouring the emergence and the persistence of polymorphism within populations are scarce. Stochastic fluctuations of population densities usually limit the levels and the duration of polymorphism in natural populations. Classical population genetics studies investigating the relative effects of genetic drift and selection regimes on the level of polymorphism [12] have highlighted that heterozygote advantage is a powerful balancing selection mechanism allowing the persistence of elevated levels of polymorphism within loci [17]. Such heterozygote advantage is frequently combined with disassortative mate preferences, whereby individuals tend to reproduce with partners displaying a phenotype different from their own. This peculiar mating behavior is promoted when heterozygous offsprings benefit from enhanced fitness, because disassortative pairs are then more likely to produce a fitter progeny. This mate preference generates powerful sexual selection promoting polymorphism within populations [19]. Disassortative mating indeed promotes rare phenotypes, because they benefit from increased mating success, therefore generating negative frequency-dependent selection.

Disassortative mating can be strict, for instance between individuals having different sexes, or between mating types as observed in fungi [2]. An emblematic example is the self-incompatibility observed in different plant families, including Brassicaceae where the *S*-locus prevents fertilization between individuals expressing the same allele [8]. In animals, disassortative behavior has also been reported, but is usually not that strict [11]. The immunity-related *MHC* locus controlling specific recognition of peptides, has also been documented to be associated with disassortative mate choice in humans [31] and mice [23], with females preferring body odours associated with *MHC* alleles different from their own. In a recent meta-analysis carried out on Primates, the propensity for *MHC*-related disassortative mating has been estimated to be relatively mild [32]. In the mimetic butterfly *Heliconius numata*, strong disassortative mating based on wing colour patterns has been documented in females, resulting in about three quarter of crosses occurring among butterflies displaying a different wing patterns in controlled experiments [6]. Almost obligate plumage based assortative mating has been reported in the white throated sparrow [29]. Altogether, disassortative mating is a rare form of mate preference, whose strength has been shown to substantially vary among animal species (see [11] for a review), highlighting the need to consider quantitative variations in the strength of disassortative preferences on the level of polymorphism maintained.

The effect of disassortative mating on the level of polymorphism has been investigated through theoretical approaches, focusing on specific examples. For instance, the polymorphism of self-incompatibility alleles within populations was specifically investigated by [33], opening many research avenues on the influence of population subdivision [24] and genetic architecture of the S-locus [3] on the number of alleles maintained at the *S*-locus, as well as on allelic turn-over [9]. The number of *S*-alleles is predicted to be large within populations, and migration of *S*-alleles among populations to be facilitated by negative FDS, therefore limiting population subdivision, consistent with empirical observations [15]. Nevertheless, the level of polymorphism maintained assuming different levels of disassortative mating is still largely unknown, and may depend on the fitness of offsprings produced by disassortative crosses.

The effect of the selection regime applied to the offsprings of disassortative crosses is also likely to have a deep influence on the amount of polymorphism maintained within populations. Offsprings of disassortative crosses are more likely to be heterozygous, and may thus benefit from increased fitness due to heterozygote advantage. But the fitness of these heterozygous offsprings may depend on the level of divergence between alleles. In the *MHC* locus for instance, empirical data provide evidence for divergent allele advantage (DDA), whereby individuals carrying the most divergent alleles benefit from increased fitness [22, 16]. Recent analyses highlight the joint effects of qualitative and quantitative variations in *MHC* alleles on fitness [1], calling for a more general view on the effect of allelic divergence on the fitness of heterozygotes.

Because the fitness advantages associated with different combinations of parental alleles may differ and ultimately determine the fate of these alleles, they are likely to interfere with the effect of the sexual selection generated by disassortative mate choice on the levels of polymorphism maintained within populations. Here we thus develop a unifying theoretical framework to explore how much the strength of disassortative preferences increases the number of alleles maintained within populations, depending on the relative fitness advantages associated with the different combinations of alleles carried by the parents.

We thus model a population with a single locus *A*, where disassortative crosses between individuals with different *A*-alleles are more successful than assortative ones. We first determine the fitness benefits associated with disassortative crosses allowing to maintain allelic polymorphism, by theoretically and numerically investigating the existence and stability (local and global) of equilibria of large dynamical systems, as well as the convergence of the population towards these equilibria. We then investigate the origin of such polymorphisms by successively introducing mutants in the population and studying the impact of such introduction on the existing diversity at the locus *A*. Finally, we consider the genetic differentiation among *A*-alleles on the persistence of polymorphism, by studying the impact of the function linking the genetic distance between parental alleles at locus *A* to reproductive advantage, therefore providing general predictions on the diversity to be expected in loci targeted by disassortative mate choice.

## 2 Level of allelic diversity maintained by disassortative mating advantage

### 2.1 General model

We consider a population of haploid individuals and a single locus *A*. Individuals reproduce sexually: they encounter mating partners uniformly at random and each mating event leads to the birth of a new offspring, with a probability that depends on the genotypes of both parents at locus *A*. We assume that crosses between individuals carrying different alleles at locus *A* (*disassortative matings*) have a greater reproductive success than crosses between individuals sharing the same *A*-allele (*assortative matings*). This may represent two different mechanisms: either individuals have disassortative sexual preferences, or the survival probability of an offspring produced by parents with different alleles at the *A* locus is higher (akin to an heterozygote advantage benefiting to the offspring).

We also assume *k* possible alleles at locus *A* in the population (denoted 1, 2, …, *k*) and no mutation. We then consider Mendelian segregation of alleles during the crosses, so that the haploid offspring inherits one allele of either parents, chosen uniformly at random.

All individuals have the same natural death rate *d*, and may also die from competition with other individuals, at a rate proportional to a competition parameter *c* > 0 and to the population density. The population is characterized at each time *t* by the respective density of individuals carrying each allele. We use an infinite population size assumption (as in [25]): we then model the dynamics of this population using a deterministic dynamical system, that can be obtained as the large population limit of a stochastic birth-and-death process, as explained in Appendix A. Let us then denote by *z_i_*(*t*) the mass represented by allele *i* in the population at time *t* 0. Then, the vector of functions (*z*_1_(*t*)*, z*_2_(*t*), …, *z_k_*(*t*))_*t*≥0_ is the unique solution of the following differential equation

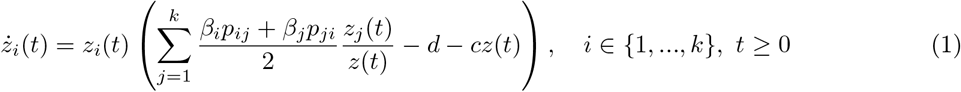

starting from 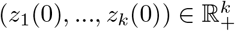, where for each *t* > 0, 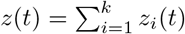 is the total population mass at time *t*. The parameter *β_i_* for *i* ∈ {1, …, *k*} is the rate at which an individual of type *i* (called first parent) reproduces, the second parent being chosen uniformly in the population. Each reproduction leads to the birth of a new individual with probability *p_ij_*, where *i* is the allele carried by the first parent and *j* the allele carried by the second parent.

We introduce

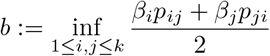

and assume that *b* > 0, implying the impossibility of having strict genetic incompatibilities between some pair of individuals. Introducing incompatibilities may however be possible and studies could be done using similar computations. For (*i, j*) ∈ {1, …, *k*}^2^, we also introduce

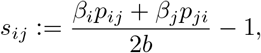

then

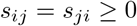

and we may rewrite (1) as

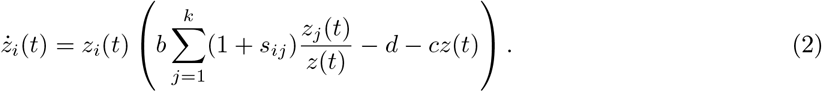

For each *i, j* ∈ {1, .., *k*} the parameter *s_ij_* may thus be interpreted as the selective advantage of a pair of parents with genotypes *i* and *j* respectively.

To maintain the population, we then always assume that

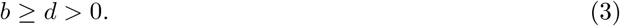

Table 1 summarizes all parameters.

**Table 1:**
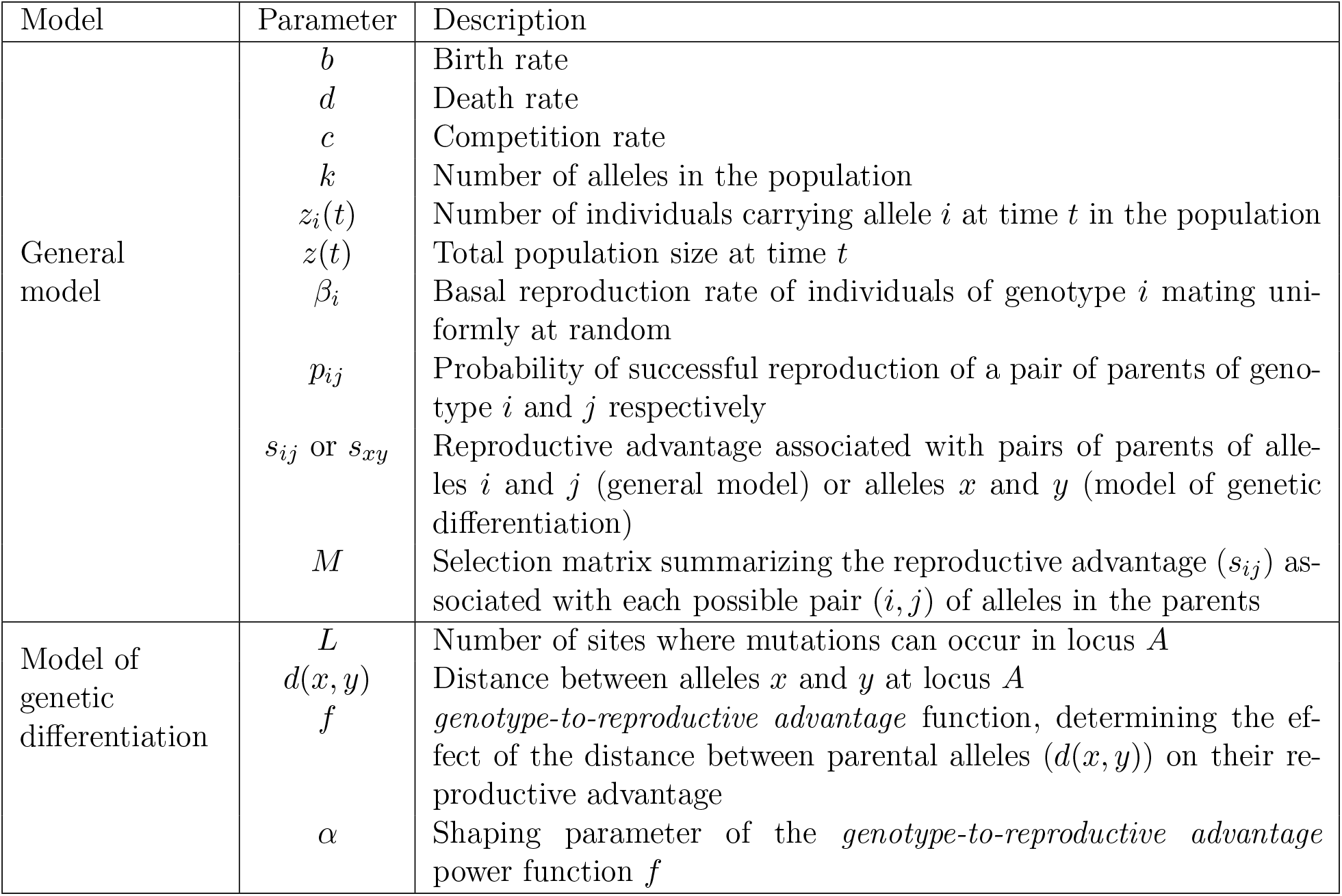
Description of parameters used in the models

### 2.2 Conditions for maintaining allelic polymorphism

In this section, we give conditions on the selective advantage of each pair of genotypes under which allelic diversity is maintained. Mathematically speaking, this diversity is preserved when the System (2) admits a positive equilibrium, and when the population converges towards this equilibrium. Our conditions depend on the matrix *M* of selective advantage:

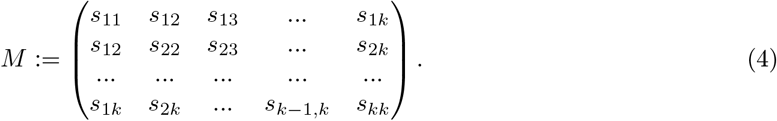

Proofs and precise mathematical results can be found in Appendix B.

**Proposition 2.1.** *Assume that* det(*M*) ≠ 0 *and*

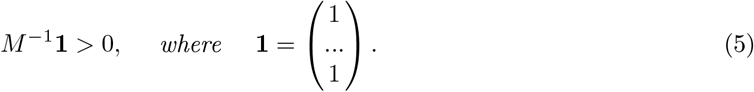

*The System* (2) *admits a unique positive equilibrium*

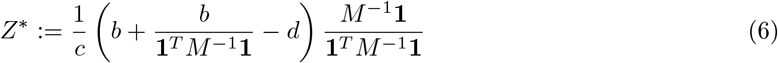

*where* **1**^*T*^ *is the transpose of vector* **1**.

*Furthermore, starting from any positive initial allelic distribution, the population will stabilize around this equilibrium if and only if the matrix M has exactly* 1 *positive eigenvalue and k* − 1 *negative eigenvalues.*

This proposition gives a condition on the selective advantage parameters *s_ij_*, under which allelic diversity will be maintained. Note that this condition depends neither on the birth rate *b*, nor on the death rate *d*, nor on the competition term *c*, but only on the disassortative advantage parameters *s_ij_*, ultimately modulating the reproductive success associated with the different allelic pairs (this is true because we assume that *b* > *d*). Note that given a matrix *M* of selective advantages, Condition (5) can be easily verified numerically. Therefore, considering a specific model for the distribution of selective advantages, one might explore how many different alleles can be maintained in the long term (see Section 3 for an example).

We then investigate more precisely this general result by studying contrasted situations, matching classical cases of overdominance. We specifically test (1) the persistence of small (*n* = 2) vs. large number (*n* ≥ 3) of alleles at locus *A*, (2) the effect of strict disassortative advantage (*i.e.* assuming *s_ij_* = 0 when *i* = *j*) and (3) the effect of equal disassortative advantages (*i.e.* assuming all *s_ij_* are equal when *i* ≠ *j*).

#### Maintaining two alleles at locus

*A* Here, the condition given in Proposition 2.1 recovers a well-documented result obtained in loci with overdominance (see [13] for instance): two alleles *A*_1_ and *A*_2_ can be maintained in a population if and only if *s*_11_ − *s*_12_ and *s*_22_ − *s*_12_ are both negative, *i.e.* when disassortative matings produce more offspring than both assortative combinations.

#### Maintaining three alleles, with strict disassortative advantage (*s_ij_* = 0 when *i* = *j*)

Let us now consider the case where three alleles can coexist at locus *A* within a population. In this case, the selection pattern is thus described by a triplet (*s*_12_, *s*_13_, *s*_23_) and the selection matrix is

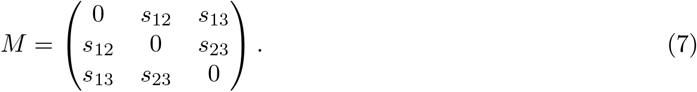

Proposition 2.1 states that the three alleles are maintained as soon as

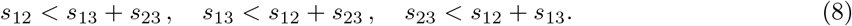

This result highlights that the three alleles are maintained when none of parental pairs has a greater advantage than the sum of the advantages of the two other possible pairs of parental alleles. This condition is for instance achieved when all disassortative pairs are similarly advantaged (*s*_12_ = *s*_13_ = *s*_23_ = *s* > 0), but can also be true for different patterns. A precise result is given in Appendix B.3.

Interestingly, this circular condition (8) was identified in [17] as a necessary condition for the maintenance of allelic diversity, in a general model of heterozygote advantage where *k* alleles are maintained. Here we demonstrate that it is not only a necessary but also a sufficient condition for the maintenance of allelic diversity in the case of three alleles.

#### Maintaining *k* alleles, when all disassortative crosses have the same reproductive advantage

We then investigate the persistence of a larger number of alleles (*k* alleles), focusing on the specific example where all disassortative pairs have the same reproductive success, *i.e.* when the matrix of interaction is given by

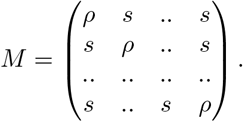

Proposition 2.1 states that allelic diversity is then maintained, as soon as *s* > *ρ* and whatever the number of alleles *k*, meaning that a large number of alleles can be maintained, when disassortative mating is favored and all disassortative pairs have equally reproductive advantages.

Our study highlights that the conditions for the persistence of a given level of allelic polymorphism at locus *A* depend on the relative reproductive advantages of disassortative *vs.* assortative crosses, but also on the relative reproductive success of the different disassortative pairs. Some conditions might allow a large number of alleles to persist, but the actual levels of polymorphism observed within population where disassortative mating is observed is also likely to crucially depend on the order of arrival of the different alleles. In particular, in cases where the reproductive advantages among disassortative pairs are not strictly equal (as in (8)), the number of alleles maintained may be strongly modified depending on the order of appearance of the different alleles.

### 2.3 Investigating the origin of polymorphism using successive introductions of mutants

To investigate the impact of the order of appearance of the mutations on the level of polymorphism, we assume that new alleles can arise in a population where one or several alleles already coexist. We thus refer to the new arising allele as a mutant and to the pre-existing alleles as resident alleles. We consider that mutations are rare enough, so that the population dynamics of the resident population reaches its equilibrium between two mutational events. We therefore aim at studying the fate of successive and non-simultaneous mutations in the population. This classical framework is close to the adaptive dynamics framework [20], because we consider rare mutation events. However, we do not assume that mutations are necessarily of small effects. We foster conditions on the mating success of the mutant allele when paired with the different resident alleles, that allow its successful invasion (*i.e.* its long-term persistence in the population).

We thus consider a population with *k* alleles, and disassortative advantage matrix is denoted by *M* as previously. We assume that *M* satisfies the conditions of Proposition 2.1, which ensure that the *k* alleles remain in the population for all times, as long as no mutation appears.

A mutant is characterized by new disassortative advantages *S^T^* = (*s*_*k*+1,1_, *s*_*k*+1,2_, …, *s*_*k*+1,*k*_) and *σ* = *s*_*k*+1,*k*+1_. We obtain (details are given in Appendix B.4) that this mutant invades if and only if

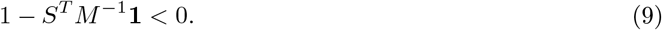

When the mutant invades the population, it can then modify the evolutionary fate of the resident alleles. We thus investigate the effect of the mutant invasion on the number of alternative alleles maintained. We are able to give a necessary and sufficient condition under which a mutant invasion leads to a population with *k* + 1 alleles maintained, *i.e.* to an increased allelic diversity after the invasion of the mutant. Let us consider the new selective matrix

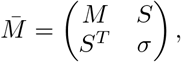

with *S* is the transpose of the line vector *S^T^*, that characterizes the population with *k* + 1 alleles. We have seen that there exists a (*k* + 1)-alleles equilibrium if 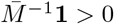. If the mutant satisfies the invasion condition (9), this is actually sufficient to ensure that this (*k* + 1)-types equilibrium is also globally asymptotically stable (see Proposition B.4 for a proof).

In the general case, we do not know the long time behavior of the population when the equilibrium with all *k* + 1 alleles does not exist; however it can be detailed in the simple case of two resident alleles or it can be studied using numerical simulations.

#### Introduction of a third allele in a resident population with two alleles

In the particular case of the introduction of a third allele in a population such that *s_ii_* = 0 for all *i*, we can determine all possible long-term behaviors of the population. Let us consider two resident alleles named 1 and 2, characterized by a disassortative advantage *s*_12_, and introduce a third allele, named 3, characterized by two new disassortative advantages *s*_13_ and *s*_23_ (the total selection matrix being then as defined in (7)). Our theoretical study leads to different cases showing that the invasion of the mutant may lead either to the coexistence of three alleles or to the extinction of one of the resident allele depending on the values of the selective advantage of the mutant, as detailed below:

- Either condition (8) holds, and the three alleles will always coexist, whatever the order of appearance of the different alleles.
- or for a given pair *i, j* ∈ {1, 2, 3}, *s_ij_* ≥ *s_ik_* + *s_kj_*. Then a further analysis of the dynamical System (2) in Proposition B.3 shows that this condition prevents the increase of polymorphism. More precisely it entails that both alleles *i* and *j* will be maintained, while allele *k* becomes extinct, regardless of whether the indices *i, j* and *k* represent the mutant or the residents.

#### Increase of polymorphism in populations where all disassortative pairs are similarly advantaged

To investigate whether high levels of polymorphism can easily be reached in populations in the favoring conditions where all disassortative pairs benefit from the same reproductive success, we study the impact of the introduction of a fourth allele in a population where three alleles are initially present and stably maintained in the population. We assume that the three resident alleles have equal interactions: *s*_12_ = *s*_13_ = *s*_23_ = *s* > 0 and that assortative mating is strictly disadvantaged, *i.e*. *s*_11_ = *s*_22_ = *s*_33_ = 0. We have seen that the three alleles coexist in such a case.

We then consider a mutant allele with (*s*_41_, *s*_42_, *s*_43_, *s*_44_) = (*x, y, z,* 0) such that the extended selection matrix is

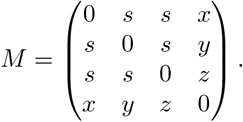

The condition for invasion of the mutant stated in (9) reads *x* + *y* + *z* > 2*s*. Depending on the values of (*x, y, z*), we investigate the number of alleles maintained in the stable equilibrium of (2) for the population. The four alleles are maintained in the population when the disassortative mating advantages associated with the different mutant/resident pairs are similar, *i.e.* when the values of *x, y* and *z* are close. When a mutant/resident pair has a strikingly higher advantage than the other pairs, then a resident allele is likely to get eliminated from the population. Interestingly, coexistence is possible even for large values of the three selective parameters *x, y* and *z* (light green area in the figure). This suggests that mutations with strong effects may invade. Highly-differentiated mutant alleles with greater disassortative advantages may then coexist with resident alleles with equal disassortative advantages. In contrast, in some cases, the equilibrium consists only in two coexisting alleles: the invasion of a new mutant may thus entail the decrease of the number of coexisting alleles.

We have given general conditions for the maintenance of allelic polymorphism within a population. This maintenance highly depends on the selective advantages associated with the different parental pairs, summarized in the matrix *M*. Interestingly, we identified some specific conditions (7) enabling the increase of polymorphism and allowing the persistence of a large number of alleles, without satisfying the restricted condition of strictly equal reproductive success in the different allelic pairs.

Because the distribution of disassortative advantages may strongly depend on the level of genetic differentiation among alleles, we then propose to study a general function linking genetic variation to selective advantages *s_ij_* and investigate the impact of this function on the emergence and polymorphism within population.

## 3 Investigating the levels of allelic differentiation maintained within population

In natural populations, the level of genetic divergence between alleles is likely to shape disassortative advantage associated with the different allelic pairs. In this second part of our study, we thus explicitly consider the effect of genetic differenciation on the selective advantages parameters, by assuming that the advantage associated with a disassortative pair is defined as an increasing function of the genetic distance between the two parental alleles. We specifically test different shapes of this function, and investigate their impact on the level of polymorphism maintained at locus *A*.

### 3.1 Modeling the link between genetic distance among alleles and their disassortative mating success

To investigate the levels of differentiation among alleles that can be maintained within population, we then consider an extension of the previous model: we assume that the allelic dissimilarity has a positive effect on the selective advantages of the different disassortative mating pairs.

In this framework, the set of possible alleles at locus *A* is {0, 1}^*L*^, where *L* is the number of sites where mutations can occur in locus *A* (fig. 2). This hypothesis is relevant to model actual loci targeted by disassortative mate choice, such as the *MHC* in vertebrates [26].

**Figure 1:**
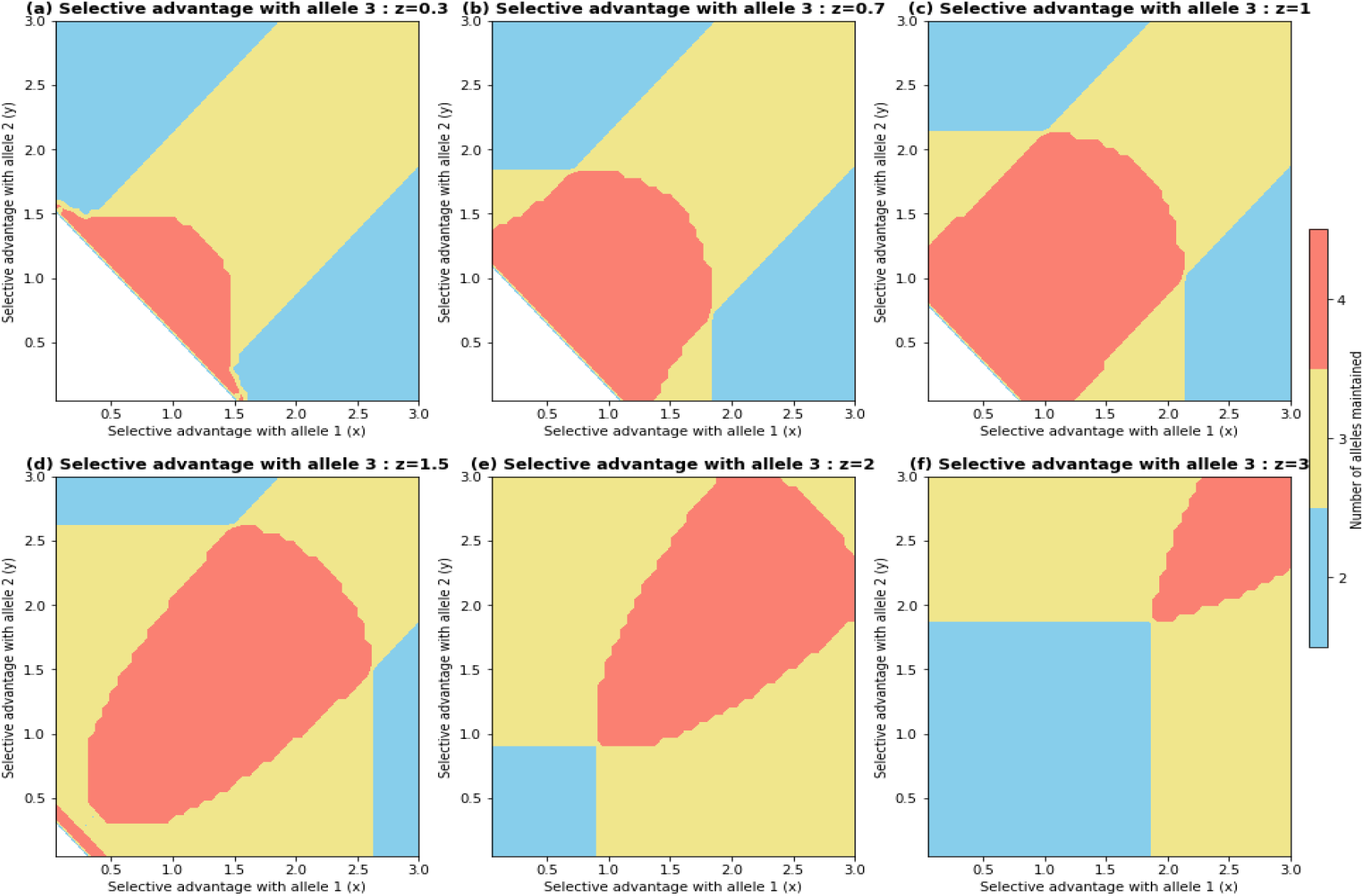
Level of polymorphism maintained in the population after the introduction of a mutant allele, depending on the selective advantage parameters associated with the pairs of parents composed of the mutant and either of the three different resident alleles (*x*, *y* and *z* respectively). In white, the mutant does not invade, in red the four alleles persist in the population, in yellow only three alleles are maintained and in blue, only two alleles are maintained. Parameters are *b* = 1, *d* = 0, *c* = 1, *s* = 1.

**Figure 2:**
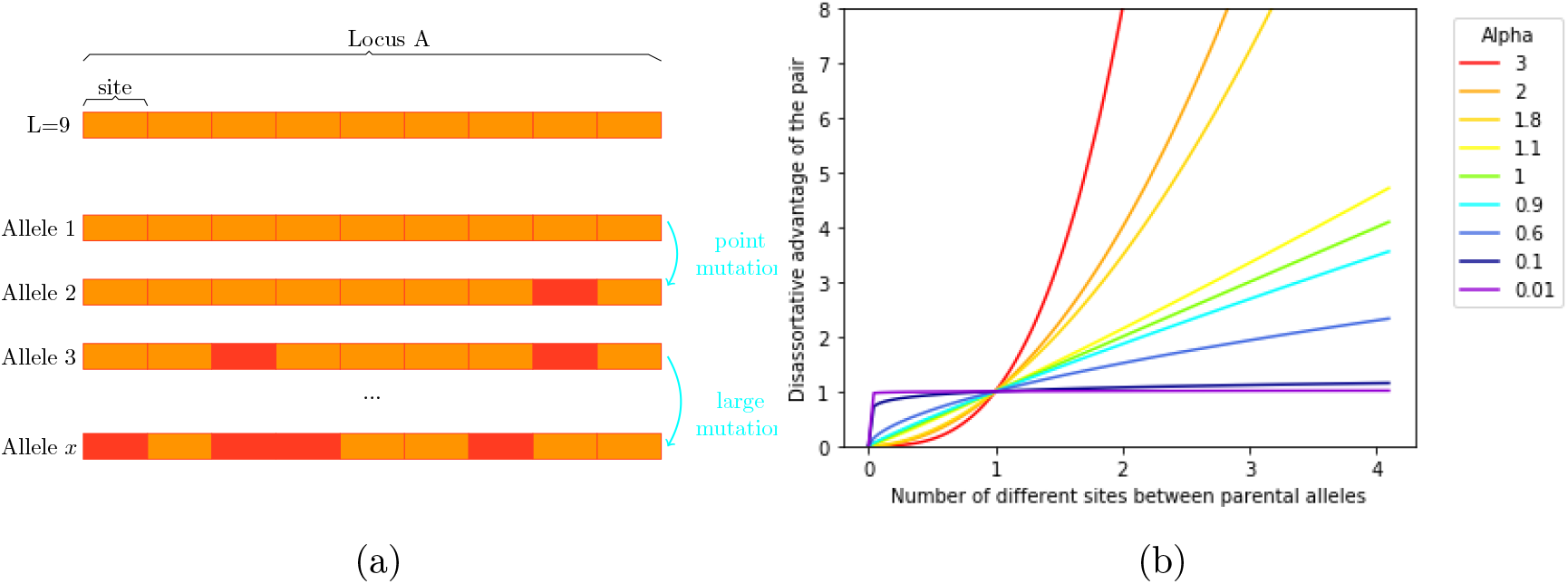
Mutation sizes and their effects on the disassortative advantage. Panel (a): The locus A contains *L* sites where mutations can occur. We model either point mutations, whereby one mutation leads to a change at a single site or other mutation kernels, where a mutation event can simultaneously affect several sites within the locus A. Panel (b): the number of sites differing between alleles in a parental pair will influence the disassortative advantage in reproduction for this pair. The parameter *α* then determines the shape of the function, *i.e.* how much the distance between alleles enhances the reproductive success of the pair. Note that in our model, we thus distinguished the size of the mutation (*i.e.* number of differing sites) from the effect of the mutations on fitness (*i.e.* the effect of genetic distance between alleles on the reproductive success).

We assume that the genetic distance between alleles carried by the parents modifies the reproductive success of disassortative pairs (fig. 2). We assume that the distance *d*(*x, y*) between two alleles *x* = (*x*_1_, …, *x_L_*) ∈ {0, 1}*L* and *y* = (*y*_1_, …, *y_L_*) ∈ {0, 1}*L* is defined by 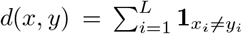. The higher the distance between *x* and *y*, the higher the reproductive success of pairs of individuals with respective alleles *x* and *y*. We introduce an increasing non-negative function *f* on ℝ_+_, such that *s_xy_* = *f* (*d*(*x, y*)), where we recall that *s_xy_* ≥ 0 is the selective advantage associated to pairs of parents with alleles *x* and *y* respectively and that it quantifies the probability for this pair of individuals to mate and produce a viable progeny. Here, we assume that the function *f* is a power function (*f* (*x*) = *x^α^* with *α* > 0). In particular the selection coefficient increases with the genetic distance, and the power *α* modulates this relationship.

We first investigate the stability of two specific final population states, chosen as the two most extreme levels of polymorphism: (1) Final population with all possible alleles maintained and (2) Final population with only the two most differentiated alleles maintained.

#### Existence and stability of a final population with all possible alleles maintained

We numerically explore the existence and stability conditions of a population state where the number of alleles maintained is 2^*L*^. Whatever the number of possible sites *L* at locus *A*, we find that the equilibrium with all possible alleles maintained in the population always exists. Indeed, each allele has the same number of alleles at distance 1, 2, …, *L* and thus the condition *M* ^−1^**1** > 0 reduces to a single condition, which is always satisfied. However, using numerical simulations (see Figure 3), we show that this equilibrium is locally and globally stable only when *α* < 1. Indeed, according to Proposition 2.1, the global stability depends on the sign of the second greatest eigenvalues of the matrix *M*, which can be computed easily numerically, even if it cannot be studied using theoretical arguments. When *α* < 1, the shape of the *genotype-to-reproductive advantage* function *f* entails that any divergent allele gains a great disassortative advantage, as soon as it slightly from the other ones, but that this advantage does not increase much when accumulating more genetic differences. This specific shape of the function may thus stabilize the polymorphism, by preventing large variations in disassortative advantages among co-existing alleles, consistent with the predictions illustrated in Fig. 1.

**Figure 3:**
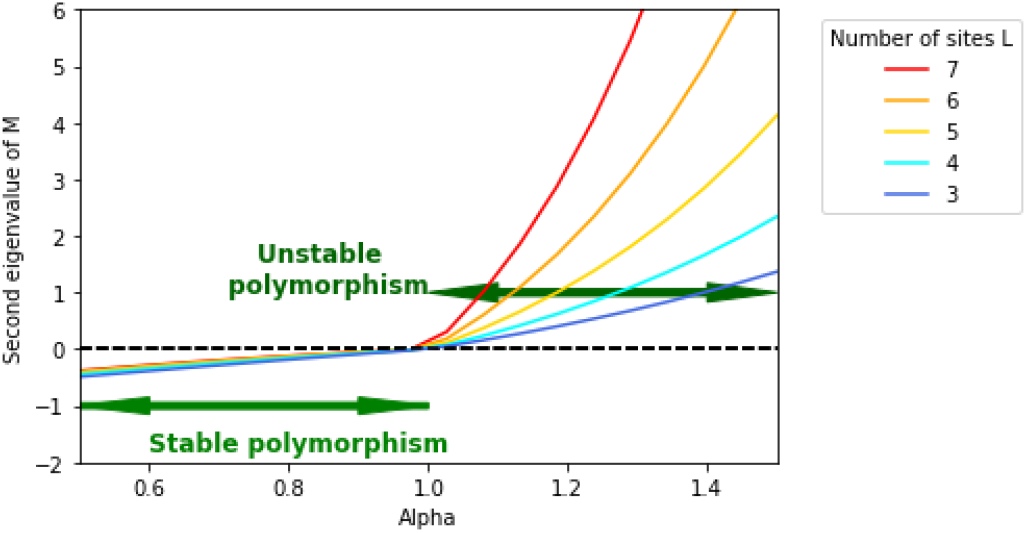
Stability of the population with all possible alleles maintained. explored using the second eigenvalue of the selection matrix *M* for different numbers of possible sites *L* at locus *A*, and depending on the shape *α* of the *genotype-to-reproductive advantage* function *f*. Note that this eigenvalue becomes positive as soon as *α* ≥ 1, therefore preventing convergence toward the polymorphic equilibrium.

#### Stability of the population state with only the two most-differentiated alleles maintained

We then explore the conditions leading to a final population composed of two alleles at maximal genetic distance *L*, *i.e. A*_1_ = (0, …, 0) and *A*_2_ = (1, …, 1). The selective advantage enjoyed in crosses between parents carrying these two alleles is *s* = *f* (*L*).

We then investigate whether a third allele might invade this population and modify the distribution of alleles. We introduce a third allele *A*_3_ ∈ {0, 1}^*L*^ \ {*A*_1_, *A*_2_}. This allele is at distance *x* from *A*_1_ and *L* − *x* from *A*_2_ for some *x* ∈ {1, · · · *L* − 1}. Therefore, the selection matrix reads

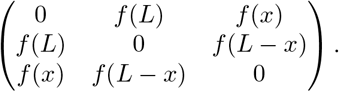

According to Proposition B.3 and using the monotony of *f*, we deduce that the invasion condition of the mutant *A*_3_ and the condition for existence and stability of a population with the three alleles can be reduced to

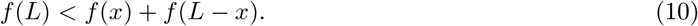

The mutant *A*_3_ will thus invade if and only if (10) holds and the final population will then be composed of the three alleles *A*_1_, *A*_2_ and *A*_3_.

In the specific case where *f* (*x*) = *x^α^*, the condition (10) is equivalent to *α* < 1. As a consequence, for *α* ≥ 1 the population with the two most differentiated alleles cannot be invaded by any new allele.

These highly-contrasted case-studies illustrate a phase transition in stability that occurs at *α* = 1, which enlightens the role of the form of the *genotype-to-reproductive advantage* function *f* in stability patterns. When *α* < 1, all alleles can be maintained simultaneously, while for *α* ≥1 a population with the two most differentiated alleles corresponds to an evolutionary stable equilibrium. When *α* < 1, the *genotype-to-reproductive advantage* function *f* has an asymptotic shape (see fig.2). This may corresponds to loci where point mutations will trigger large disassortative advantage, while highly differentiated variants have little advantages as compared to slightly divergent ones. This is likely to promote the emergence of a large diversity of alleles with similar levels of disassortative advantages. On the contrary, when *α* > 1, each mutation accumulating in the locus *A* induces a supplementary advantage and might therefore replace less differentiated resident alleles.

#### Diversity of allelic distribution

The analytical study of other cases of persistence of any given sample of the 2^*L*^ different alleles is highly challenging. However, using our criteria, we numerically explore the conditions of emergence of populations with a large number of alleles. When several alleles are maintained within populations, we can then explore the number of individuals carrying each of the different alleles maintained in the population. (See Figures 4).

**Figure 4:**
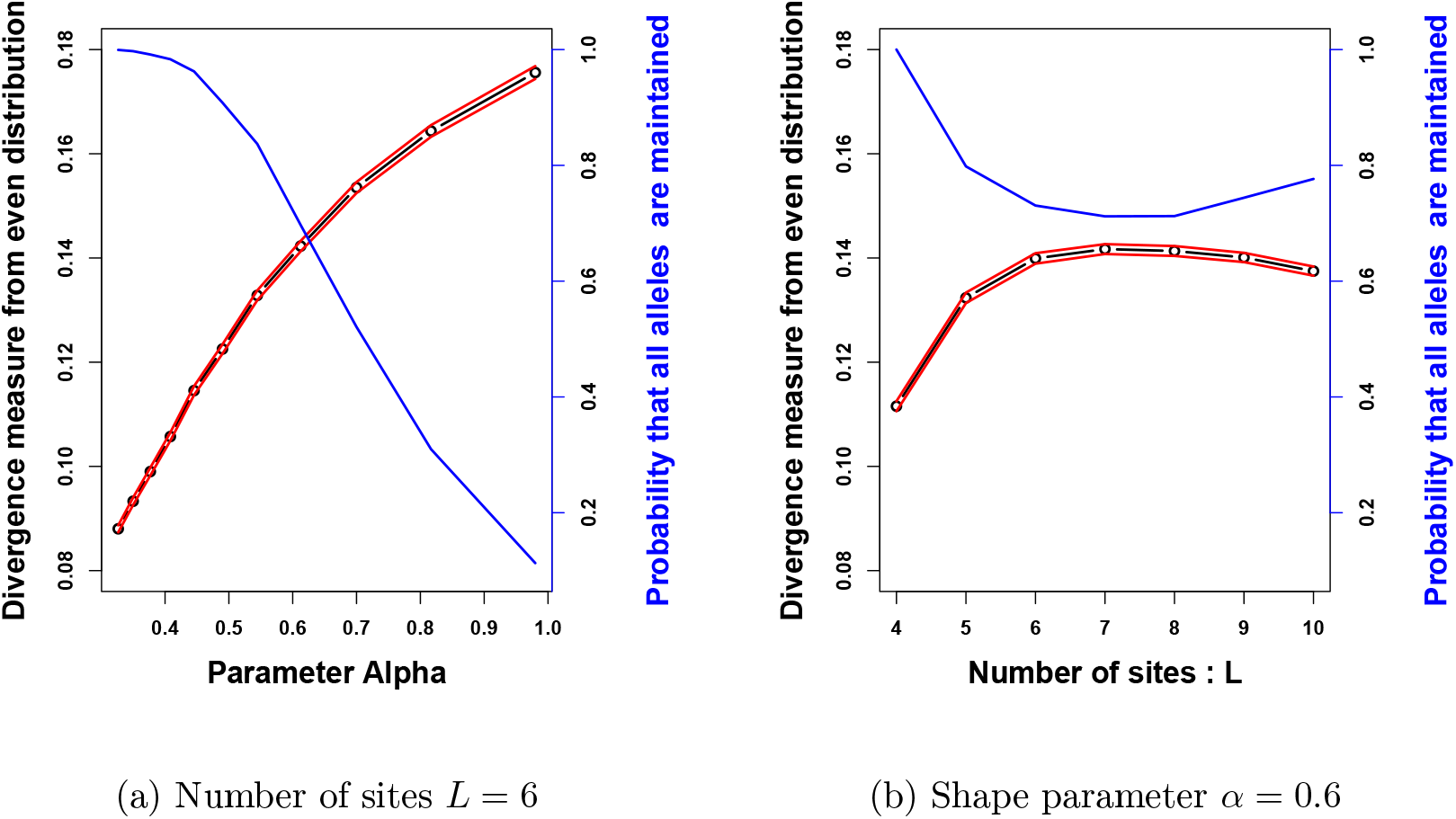
Probability of persistence of 10 randomly chosen alleles in the population (in blue) and departure in the actual distribution of these alleles in the population, from an even distribution of alleles (in black) for different values of the number of sites within locus (*L*) and of the parameter *α*, tuning the shape of the *genotype-to-reproductive advantage* function. The red lines corresponds to the 95% confidence intervals computed over the stochastic simulations. On panel (a), the number of sites *L* is fixed to 6. The distribution departs further away from an uniform distribution when *α* increases, while the probability of maintaining 10 alleles drops. On panel (b), the shape parameter *α* is fixed to 0.6. The departure from even distribution first increases with the number of sites, and then decreases when the number of sites is high. The probability of choosing 10 alleles that can be maintained decreases slowly when the number of sites increases.

To this aim, we draw uniformly at random populations of 10 alleles among the 2^*L*^ possible alleles, we then check whether these alleles can coexist. This stochastic procedure was repeated in order to obtain 5000 possible populations and allows to estimate the probability that 10 randomly chosen alleles are maintained in the population, that is when the associated matrix of selective advantage satisfies (5). Note that for the rest of the study, we thus only consider the populations in which the 10 alleles are maintained. For each population of 10 alleles obtained, we then compute the number of individuals carrying each allele. Finally, we measure the difference between these numbers and the numbers obtained when alleles are equally distributed in the population. Figure 4 shows the empirical mean of this measure for a large number of replicates, for different values of *L* and *α* ≤ 1, as well as the probability of maintaining 10 alleles chosen uniformly at random within the population.

On Figure 4a, the number of sites *L* is fixed to 6 such that we select randomly 10 alleles among the 2^6^ = 64. We observe that the probability of choosing 10 alleles that will be stably maintained in the population drops when *α* increases. This is consistent with previous conclusions stating that for *α* ≥ 1 only small numbers of alleles can stably persist in the population. Furthermore, we notice that in the population where ten alleles are stably maintained in the populations, the distribution of allele frequencies departs further from a uniform distribution, when *α* increases. Larger values of *α* are indeed likely to increase variation in the reproductive success of the different allelic pairs. Fr a given set of alleles, their associated selective advantages indeed depart further away from the constant case as *α* increases (see fig. 2).

On Figure 4b, the shape parameter *α* is fixed to 0.6 and we modify the number *L* of sites and therefore the number of possible alleles 2^*L*^. We observe that the departure from even distribution first increases with the number of sites (up to *L* = 7), and then slightly decreases when the number of sites is high. The initial growth in the departure from the even distribution may stem from the increased number of possible alleles 2^*L*^, enhancing the diversity in the selected alleles. Nevertheless, when *L* is large enough, the increase in the number of possible alleles no longer impacts the diversity in the chosen alleles because the selected alleles are then very likely to be quite differentiated. When *L* is getting higher, the matrix of selective advantages may then approach a matrix with equal disassortative advantages. Note that for large values of *L* the choice of 10 alleles among a large panel will create very diverse selection patterns strongly differing from the simple case where *s_ij_* = *s* for all pairs. This then leads to population structure with various allele frequencies.

### 3.2 Emergence of allelic diversity

We aim at exploring how allelic diversity might emerge from multiple rounds of mutant invasions. We first assume that each mutation affects only one site within locus *A*, which is then shifted towards the opposite value. When an offspring is born, it is either similar to one of its parents, or it has only one site different from one of its parental alleles. The mutant allele is then at a distance one from this parental allele. We observe two different evolutionary outcomes depending on the shape of the *genotype-to-reproductive advantage* function *f*, determined by the value of *α*, as detailed below.

#### 3.2.1 *Genotype-to-reproductive advantage* function where disassortative advantage saturates when differentiation between allele increases (*α* < 1)

From our theoretical study described above, we recall that:

1. Any resident population with two alleles *A*_1_ and *A*_2_ can be invaded by any new mutant *B* at distance 1 of either *A*_1_ or *A*_2_ and will lead to a population with 3 alleles.
2. The population with all possible alleles maintained then exists and is stable.

However, we have no theoretical evidence that the successive introductions of mutants will lead to the population maintaining a large number of alleles at distant 1 from each other. We thus numerically explore the successive invasions of mutations, using simulations encoded in Python.

Figure 5 illustrates the contrasted evolutionary fates followed by the introduced mutants: the mutant can either go extinct rapidly if its fitness is negative, or it can invade the population. If it invades, it either co-exists with all resident alleles or triggers the loss of one or several resident alleles. The right panel (Fig. 5a) explicitly shows the allelic turn-over through time, suggesting that the number of alleles can be high but may also strongly vary through time in natural populations. Note that the different alleles do not have the same frequency in the population, even when the number of alleles maintained is large.

**Figure 5:**
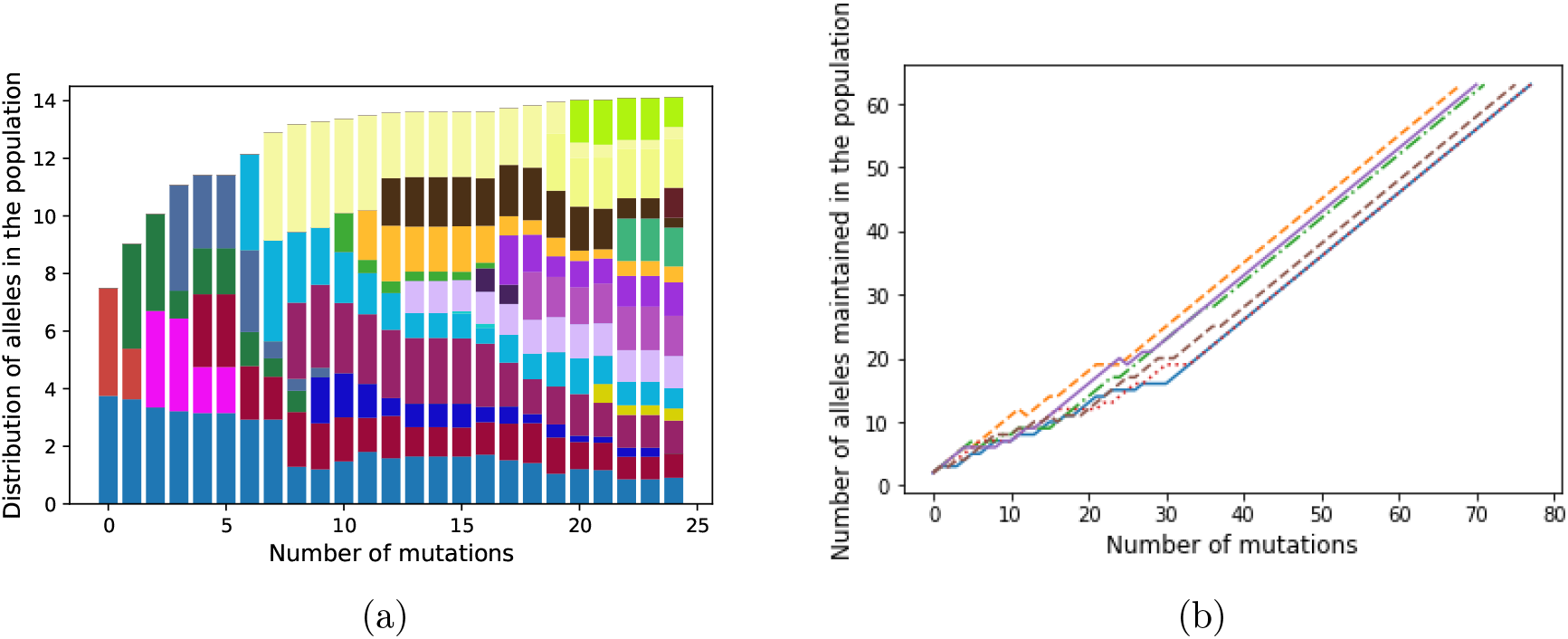
Evolution of the number of alleles maintained in the population, assuming point mutations and convex shape of function determining the fitness of allelic pairs (*α* ≤ 1). From an initial population with two alleles, we numerically induce successive mutations and track down their invasion success through time. Panel (a) shows the distribution of alleles in the population through time. Each color corresponds to a given allele and the height of the bar is the number of individuals carrying each allele within the population at a each time. Panel (b) gives number of alleles maintained at equilibrium after each mutation until the total number of alleles is reached. Each line corresponds to a different numerical simulation (*n* = 6). Here *L* = 6 and *α* = 0.6 such that there are 2^6^ = 64 possible alleles.

Figure 5b then highlights that, after a sufficiently large number of mutations, all possible alleles are maintained in the population. We furthermore observe that when the number of coexisting alleles reaches a sufficiently high level, any new mutant invades and increases the allelic diversity.

#### 3.2.2 *Genotype-to-reproductive advantage* function where disassortative advantage is always enhanced when differentiation between allele increases (*α* ≥ 1)

. We prove that in this case, only two alleles can coexist in the population through time, meaning that any mutant invasion leads to the extinction of one out of the two resident alleles. We assume that the initial population is composed of two alleles *A*_1_ and *A*_2_ at distance *x*. When a mutant *A*_3_ arises, it is at distance 1 of its parental allele (say *A*_1_ by symmetry) and at distance *x* + 1 or *x* − 1 of the other parental allele *A*_2_. This particular property arises from the choice of the mutation kernel (see the section below on the influence of the mutation kernel).

When assuming that the mutant is at distance *x* − 1 of *A*_2_, the invasion condition reads 1 + (*x* − 1)^*α*^ − *x^α^* > 0, and is thus never true for *α* ≥ 1. Therefore, a mutant allele closer from the resident allele *A*_2_ than the resident allele *A*_1_, can never invade the population. In contrast, when assuming that the mutant is at distance *x* + 1 of *A*_2_, then the invasion condition reads 1 + (*x* + 1)^*α*^ − *x^α^* > 0 and is true, since *x* ↦ *x^α^* is increasing for *α* > 0. By applying Proposition B.3, we can deduce that, when the mutant allele invades, the allele *A*_1_ is eliminated from the population. The resulting population is then composed of the two most differentiated alleles *A*_2_ and *A*_3_, at distance *x* + 1. In case of successive emergence of new alleles by point mutations, we thus observe increasing distances between the pairs of alleles maintained in the evolving population. This result goes further than the global stability of the population formed of the two most differentiated alleles: it proves that starting from any initial couple of alleles, the successive mutations always increase the genetic distances between the two surviving alleles. As a consequence, after a sufficient number of mutations, the population will be composed of two alleles at distance *L*. However, this general fate of the populations can be modified when mutations affecting several sites together arise.

##### Importance of the mutation kernel

We then investigate the effect of the number of sites within the locus *A* affected by a mutational event. In this section, we develop an example showing that, with this kind of mutation kernels, coexistence of more than two alleles can be observed, even if *α* ≥ 1.

We now assume that the mutants are chosen uniformly at random among the possible alleles and that these new mutants are introduced successively in the population. These introductions are repeated until we obtain a set of alleles which cannot be invaded by any new mutant allele. The following result gives the law of the final number of co-existing alleles in the population for a particular example ( *i.e.* assuming *L* = 3 and *α* ≥ 1). In particular, we obtain from a mathematical reasoning that the number of alleles maintained is not always equal to 2 and depends on the order of mutant introductions.

##### Proposition 3.1

*If α* [1, 2] *and L* = 3 *then the final number of co-existing alleles is equal to* 2 *(with probability* 51/56), *or to* 4 *(with probability* 5/56).

This proposition illustrates the importance of the mutation kernel on the final number of alleles and on the distribution of their genetic distances. The proof of Proposition 3.1 is given in Appendix C.2.

We then use numerical simulations to observe the evolution of the number of coexisting alleles for *α* ≥ 1, when the number of sites *L* at locus *A* increases. We use a Monte Carlo procedure: we repeated 5000 times the evolution of the population, for uniform mutations on all the possible alleles at locus *A*, until we obtain a stable population. This allowed us to obtain a vector containing the number of alleles maintained in the final population within each repetition. Figure 7 represents the histogram obtained for different values of *L* and *α* ≥ 1. Consistent with the theoretical results obtained above for *L* = 3, that for *α* ≥ 2 the limiting population is always composed of 2 alleles, while for 1 ≤ *α* < 2 it can be composed of 4 alleles (Figure 7a). A similar pattern is observed for *L* = 4 on figure 7b. Larger number of alleles are maintained from the successive introductions of uniform mutations when *L* = 5 or *L* = 6 (see Figure 7c and 7d). These populations with larger numbers of alleles arise with a small probability (of order 10^−3^) and are more frequent for small values of *α*. This highlights that stable co-existence of more than 2 alleles within population exists even in the case where *α* > 1. They arise more frequently for large values of *L*, possibly because the number of allelic combinations fitting the coexistence condition increases.

## 4 Discussion

### 4.1 A general model predicting balanced as well as unbalanced polymorphism in loci under disassortative and/or heterozgote advantage

In this article, we explored the conditions of emergence and persistence of allelic diversity at a locus where disassortative pairs of parents benefit from increased reproductive success. Our haploid model can therefore cover the cases explored in models exploring persistence of polymorphism in either (1) loci where heterozygote advantage occurs [17] or (2) loci where disassortative mating happens, such as the self-incompatibility locus in plants [33]. From our general model, we retrieve classical conditions of maintenance of large number of alleles at these loci: we indeed confirm that a large number of alleles can be maintained within population when the advantages associated with the different dissassortative pairs (akin to the different heterozygote advantages in diploid models exploring overdominance) are close to each other. Our disassortative advantage *s_ij_* corresponds to the fitness of the genotype *A_i_A_j_* denoted by *W_ij_* in [17]. However, the main difference is that we do not consider the dynamics of allele frequencies within a population with fixed size, but instead, we modeled the dynamics of both the alleles and the total population size. In our case, the dynamics of allele frequencies solves:

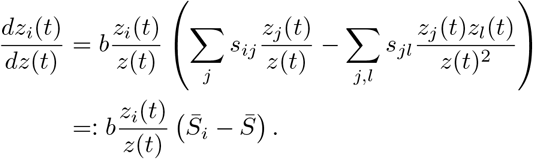

As a consequence, if an equilibrium exists, it will satisfy 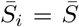 similarly as Equation (2) in [17]. We then provide the sufficient and necessary conditions for the convergence of solutions of the dynamical system to the equilibria. Our mathematical analyses thus reinforce the necessary conditions for existence and local stability obtained in [17].

In these classical models where the advantages associated with all allelic pairs are equal, even distribution of the frequencies is predicted. However, here we observed equilibria with unbalanced frequencies within populations, depending on the genetic distances among alleles and their respective effect on the fitness of the different pairs. Such uneven frequencies are frequently observed in natural populations at the self-incompatibility locus in plants [18]. The effect of the dominance relationships among alleles at the *S*-locus has for instance been documented as a major effect tuning allelic frequencies in loci under disassortative mating [30]. Here we propose a general framework allowing to investigate the effect of (1) each mutational event on the genetic distance between alleles as well as (2) the function linking genetic distance to fitness advantage associated with each allelic pair (Section 3.1). This general framework provides general expectation on the level of polymorphism, the evolution of the allelic divergence as well as on the allelic turnover.

### 4.2 Allelic diversity and turn-over

By explicitly considering the successive introductions of mutants (Section 2.3), our model provides general predictions on the emergence of genetic diversity in the population. We obtain a simple criterion on the disassortative mating parameters, under which a new mutant can invade. The final genetic diversity of the population is however difficult to determine from the general model. We nevertheless obtained a simple matricial criterion enabling the co-existence of both the mutant and all resident alleles. Besides, the behavior of the population with two resident alleles after the introduction of a third allele was also fully characterized in the general model.

When considering explicitly the genetic distance among alleles and its effect on the fitness of disassortative pairs (Section 3.1), we highlight the strong effect of the number of sites affected by a single mutational event. The genetic diversity maintained within population is maximal when only mutations between close genotypes are possible. Such mutational events may correspond to point mutation of small phenotypic effect appearing by *de novo* within natural populations. However, in loci under disassortative mating or heterozygotes advantage, the migration of alleles from different populations [15], as well as the introgression [4] from closely-related species is frequently observed. Such immigrants or introgressed alleles are likely to be quite different from the resident alleles and to unbalance the populations of resident alleles and limit the polymorphism maintained. In natural populations, variants with few genetic differences are shown to co-exist with highly differentiated variants stemming from migration between populations or species [5], therefore complexifying the dynamics of polymorphism predicted by our model.

### 4.3 Allelic differentation

By explicitly testing the effect of the shape of the function linking genetic distances among alleles to associated disassortative advantages (Section 3.1), we show the significance of the shape of the *genotype-to-reproductive advantage* function on both the number of alleles maintained and their level of genetic divergence.

Our model indeed shows that, when disassortative advantage only poorly increases when allelic differentiation becomes large, then great number of alleles can be maintained within population, with substantial turn-over among alleles. This may correspond to the situations observed at several mating types loci or at the self-incompatibility locus or as the MHC loci, where large number of alleles are maintained. At these loci, few mutations already have a significant functional effect, thus promoting their maintenance. On the contrary, when the reproductive advantage still increases when differentiation levels between alleles become larger, few highly differentiated alleles are predicted to be maintained within the population. This would correspond to the large differences observed between haplotypes at the supergene controlling color patterns submitted to disassortative mating in the butterfly *H. numata* [10] or in the loci controlling plumage coloration in the *Z. albicollis* sparrow [28]. While other shapes of the *genotype-to-reproductive advantage* function may exist, our model still offers a mechanistic explanation for strikingly different levels of polymorphism and allelic differentiation observed in different mating systems.

We now hope that our theoretical predictions will simulate empirical efforts in characterizing the respective fitness advantages associated with the different heterozygotes or disassortative crosses, enabling to better understand the evolution of differentiation among alleles in these systems.

### Numerical simulations

The code for this article is available at https://plmlab.math.cnrs.fr/costa2150/heterozygote_advantage

## Acknowledgements

This work has been supported by the Chair “Modélisation Mathématique et Biodiversité” of Veolia Environnement-École Polytechnique-Muséum national d’Histoire naturelle-Fondation X. This work has been partially supported by the LabEx PERSYVAL-Lab (ANR-11-LABX-0025-01) funded by the French program Investissements d’avenir

We warmly thank Clément, Gabriel and Lucie for enlarging our world during this work.

## A Birth-and-death model and convergence to dynamical system

The dynamical system (1) governing the population dynamics can be obtained as a large population limit of a stochastic individual based model (similarly as in [7, 25] for instance).

More precisely, let *K* be a large parameter that gives the order size of the population. The microscopic population is represented by the process 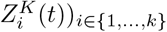 where 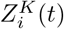 gives the size, divided by *K*, of the population of type *i* individuals at time *t*. Each individual of type *i* reproduces with rate *β_i_*, that is to say after a time distributed as an exponential random variable of parameter *β_i_*. As explained in the main text, the second parent is chosen uniformly at random among all individuals alive in the population. If the second parent is of type *j*, with probability *p_ij_* this mating event leads to the birth of a new individual, being of type *i* with probability one half, and of type *j* with probability one half when there is no mutation (otherwise the mutation is applied to the genotype *i* or *j* with the same probability). All the indivivuals have the same death rate, which is given by

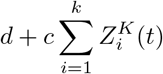

at time *t*. Under the assumption that the sequence of initial conditions 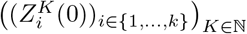 converges (in probability) when *K* goes to infinity, the sequence of stochastic functions 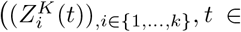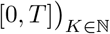 also converges (in probability for the uniform convergence) to the trajectory of the deterministic system (1).

## B Proof of the main results

This section gathers the theoretical results on the behavior of the solutions to system (2):

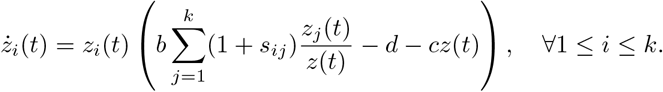

Recall the matrix *M* = (*s_ij_*)_1≤*i,j,*≤*k*_

### B.1 Existence of positive equilibria

Our first result states conditions for a positive equilibrium to exist, meaning that each coordinate of the vector is strictly positive.

#### Proposition B.1

*If* det(*M*) ≠ 0, *the system admits a positive equilibrium if and only if M* ^−1^**1** > 0. *In this case, the equilibrium is unique and is given by*

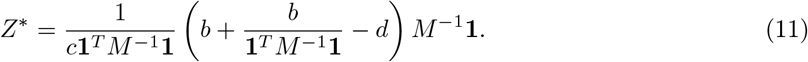

*If* det(*M*) = 0, *the system may have no equilibrium or a infinite number of equilibria, whose space corresponds to the intersection of an affine space of direction the kernel of M with the cone of positive coordinates*.

*Proof of Proposition B.1.*

<u>**Case** det(*M*) ≠ 0:</u> Firstly, we assume that there exists a positive equilibrium 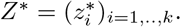. From (2), the following equality holds for every *i* ∈ {1, .., *k*},

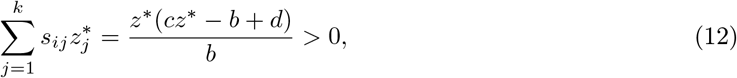

with 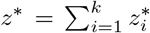. Note that this equation implies that *cz*^∗^ > *b* − *d*, since *s_ij_* > 0. Using a matrix formulation, we have the following linear system:

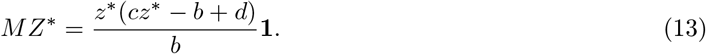

As det(*M*) ≠ 0, the matrix *M* has an inverse and

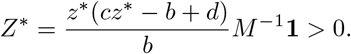

Since 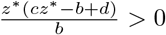, we thus deduce that *M* ^−1^**1** > 0.

Moreover by summing all coordinates, we find

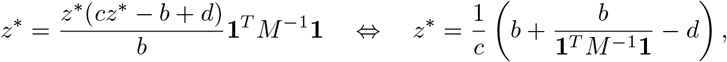

and finally, the equilibrium is unique and defined by (6).

On the other way, if we assume that *M* ^−1^**1** > 0, then we can define *Z*^∗^ by (6) and, using the above computation, it is straightforward to verify that it is a positive equilibrium of (2). This ends the proof for the case det(*M*) ≠ 0.

<u>**Case** det(*M*) = 0:</u>

Let us verify whether there exists a vector *X* with positive coordinates such that *MX* = **1**.

Indeed, if such a vector *X* does not exist, then (12) can never be satisfied and there is no equilibrium to (2).

On the contrary, if we can find such a vector *X*, we set

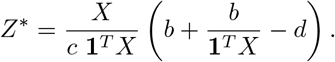

Then it is straightforward to verify that 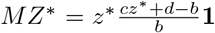, with *z*^∗^ = **1**^*T*^ *Z*\*. Hence, *Z*^∗^ satisfies (12) and is a positive equilibrium to (2). However, this equilibrium is not unique. Indeed, **1** belongs to the image of *M*, and since *M* is a symmetric matrix, any vector *Y* in the kernel of *M* is orthogonal to **1**, i.e. **1**^*T*^*Y* = 0. Then, for all vector *Y* ∈ *Ker*(*M*), 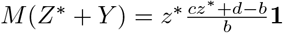, with *z*^∗^ = **1**^*T*^ *Z*^∗^ = **1**^*T*^ (*Z*^∗^ + *Y*), and *Z*^∗^ + *Y* is also an equilibrium to (2).

We now construct examples for the two possibilities. Assume that *M* can be written as

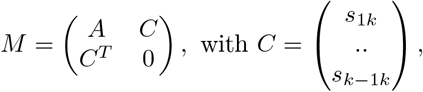

det(*A*) ≠ 0 and such that *A*^−1^**1** > 0. Since *A* is a symmetric matrix, there exists an orthonormal basis (*V_i_*)_*i*=1,..,*k*−1_ of ℝ^*k*−1^ where *V_i_* is an eigenvector of *A* associated to its non-null eigenvalue *λ_i_* for all *i* ∈ {1, .., *k* − 1}. Let us decompose *C* in this basis 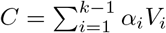.

Our aim is to construct *C*, and thus to select (*α_i_*)_*i*=1,..,*k*−1_, to find the examples. First, we have to select (*α_i_*)_*i*=1,..,*k*−1_ such that 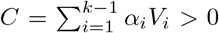. Then using the Schur complement, we have that det(*M*) = −(*C^T^ A*^−1^*C*) × det(*A*) and thus det(*M*) = 0 if and only if

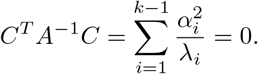

This gives us a way to construct a matrix *M* with det(*M*) = 0.

Finally, to ensure that there exists *Y* > 0 with *MY* = **1**. Let us consider vectors depending on a parameter *η* ∈ ℝ such that

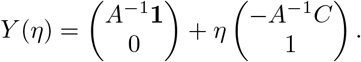

Let us study the image by *M* of this vector

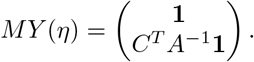

Thus if 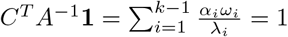 where 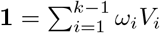, then we can find *η* > 0 sufficiently small, such that *MY* (*η*) = **1** with *Y* (*η*) > 0.

With this in mind, we can explicit an example for each possibility. Let

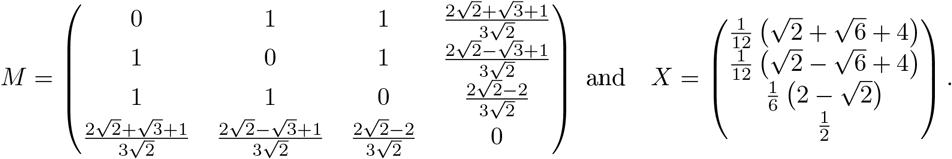

*M* and *X* have non-negative coordinates, det(*M*) = 0 and *MX* = **1**. Thus, in this example, the associated system of equations (2) has an infinite number of positive stationary states.

Now, let *a* > 0 and

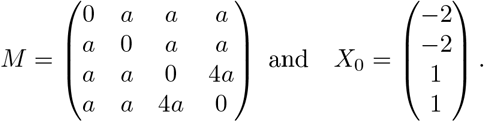

In this example, det(*M*) = 0, *Ker*(*M*) = *V ect*(*X*_0_) = *Im*(*M*)^*T*^, however **1**^*T*^ *X*_0_ ≠ 0 hence **1** does not belong to the image of *M*. In other words, the associated system of equations (2) has no positive stationary state.

#### B.2 Convergence toward the positive equilibria

We now aim at describing the dynamics of the population described by system (2). The following proposition gives conditions for a positive equilibrium to be stable. Moreover, it states that conditions for local stability and global stability are identical, *i.e.* a locally stable equilibrium attracts all trajectories starting with positive initial conditions.

##### Proposition B.2

*Assume that* det(*M*) ≠ 0 *and M* ^−1^**1** > 0, *and denote the unique positive equilibrium to* (2) *by Z*^∗^, *whose expression is given in* (6).

*i) Z*^∗^ *is locally stable if and only if M has* 1 *positive eigenvalue and k* − 1 *negative eigenvalues*.

*ii) Z*^∗^ *is locally stable if and only if it is globally stable on* 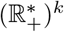.

Our proof relies on a study of the Jacobian matrix of the system computed at the positive equilibria and on the construction of a Lyapunov function. Actually, we prove that the criterion on the eigenvalues of *M* is equivalent to the fact that the Jacobian matrix of the system admits only negative eigenvalues. The second point of the proposition will be proved using the following function

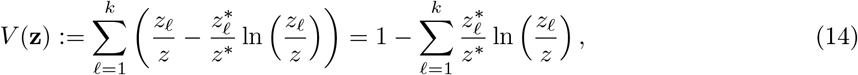

which is a Lyapunov function for the dynamical system (2).

*Proof.* <u>Proof of *i*):</u> Let us first give the Jacobian matrix of the system (2) around equilibrium *Z*^∗^. For all *i* ∈ {1, .., *k*}, let

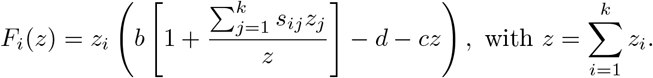

Recall that 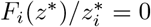. Then by differentiating the function *F_i_* w.r.t *z_i_* and *z_j_* for *j* ≠ *i*, we find

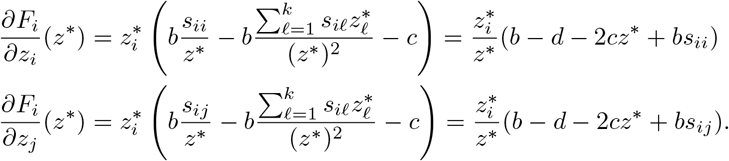

Hence the jacobian Matrix of the system can be written

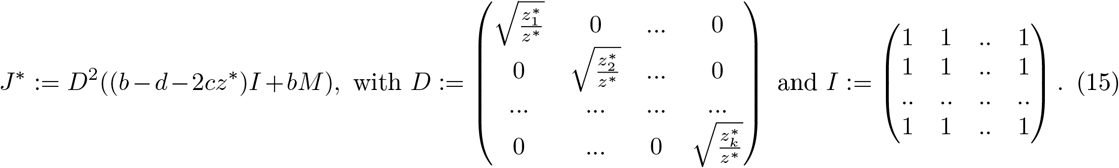

Then since for any 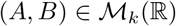, det(*AB*) = det(*BA*), we have

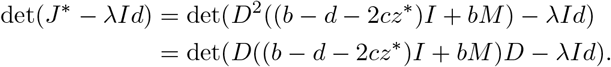

In other words, the eigenvalues of *J* ^∗^ are the same as the ones of

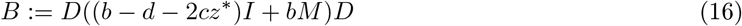

which is symmetric and has real eigenvalues. According to Proposition 2.1,

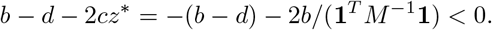

Hence, we can write the matrix *B* as

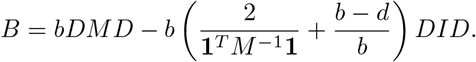

Let us prove that *B* has negative eigenvalues if *M* has only 1 positive eigenvalue. To this aim, notice that as *M* and *DMD* are congruent, according to Sylvester’s law of inertia [27], they have the same numbers of positive/negative eigenvalues.

Let us first assume that *M* as well as *DMD* have only 1 positive eigenvalue. As *DMD* is symmetric, there exists an orthonormal basis (*V_i_*)_*i*=1,..,*k*_ of ℝ^*k*^ formed with eigenvectors of *DMD*, associated to its eigenvalues (*λ_i_*)_*i*=1,..,*k*_. In the sequel we denote by *λ*_1_ > 0 the positive eigenvalue. Using the definition of *D* in (15), we obtain that

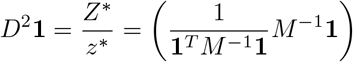

combining (13) and (6). Then

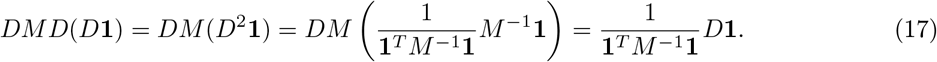

Since by assumption 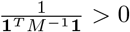, we deduce that it is the unique positive eigenvalue of *DMD* and that *D***1** is a positive eigenvector associated to the positive eigenvalue of *DMD*.

From (17), we have

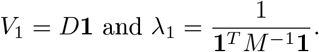

Now, let us consider any *Y* ∈ ℝ*k* and its decomposition in the eigenvectors’ basis: there exist (*γ*_1_, .., *γ_k_*) ∈ ℝ^*k*^ such that 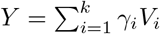, and

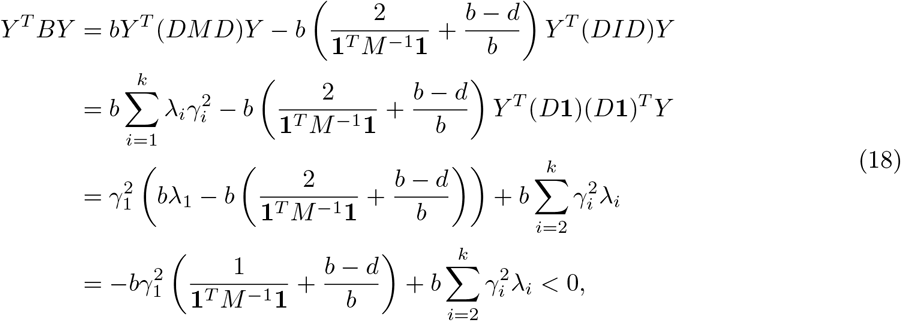

since the eigenvalues *λ*_2_, .., *λ_k_* of *DMD* are negative by assumption. In other words, *B* is a symmetric definite negative matrix. Recalling that *B* and the jacobian matrix *J* ^∗^ have the same eigenvalues, we deduce that the equilibrium is locally stable.

Conversely, if *M* has more than 1 positive eigenvalue, it is also the case of *DMD*. Then, assuming for example that *λ*_2_ > 0, we have, by an application of (18) with *Y* = *V*_2_,

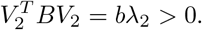

Hence *B* has at least 1 positive eigenvalue. The stationary state is unstable.

<u>Proof of *ii*):</u> We are now ready to prove the second point of the proposition. We only have to prove the direct implication, as the other one is obvious. To this aim, let us assume that the equilibrium is locally stable and thus all eigenvalues of *J* ^∗^ are negative. Recall the definition of the function *V* in (14). By differentiating, we find

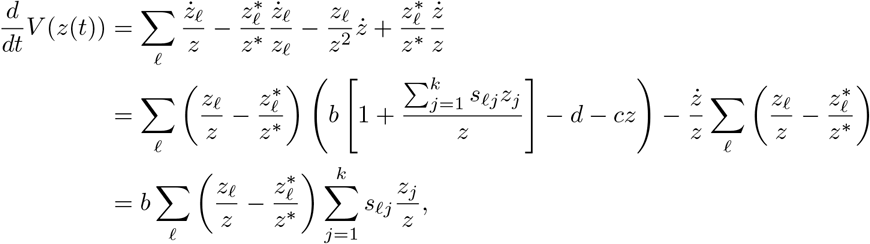

since 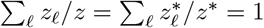. In addition with the fact that 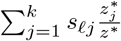 is a constant, we find

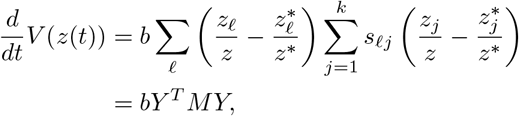

with

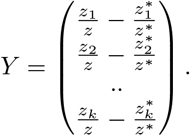

Now, we recall that the stationary state is assumed to be locally stable, i.e. all eigenvalues of *J* ^∗^ (or *B*) are negative. Since *B* and (*b* − *d* − 2*cz*^∗^)*I* + *bM* are congruent, these matrices are both symmetric definite negative. Furthermore, for all *X* ∈ ℝ^*k*^ \ {0}, *X^T^* ((*b* − *d* − 2*cz*^∗^)*I* + *bM*)*X* < 0. Since **1**^*T*^ *Y* = 0, we find

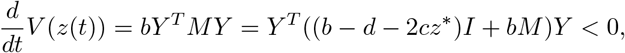

as soon as *Y* ≠ 0, and 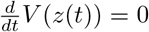 if and only if *Y* = 0, i.e. 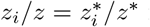 for all 1 ≤ *i* ≤ *k*. Moreover, we can prove that the set {*z* ∈ ℝ^*k*^, *Y* = 0} is a positive invariant set for the dynamical system (2). Indeed, for all *i* ∈ {1, .., *k*},

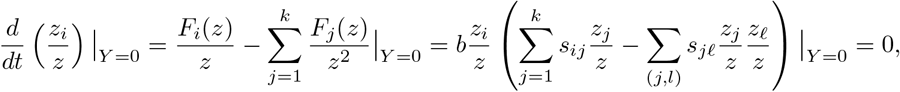

where we used (12). Theorem 1 of [14] then ensures that any bounded positive solution to (2) converges toward this invariant set. Finally, we prove that the solutions of (2) remain bounded by studying the dynamics of the total population size *z*, which satisfies

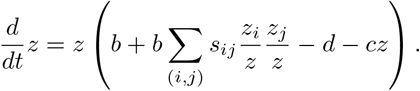

We easily obtain that

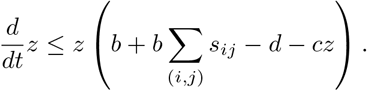

and thus classical results on logistic equation ensure that the total population size remains bounded through time.

Lasalle Theorem then ensures that 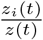 converges to 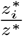. To conclude the proof it remains to prove that *z*(*t*) converges to *z*^∗^. Let us write 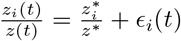 and according to (12)

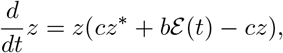

with

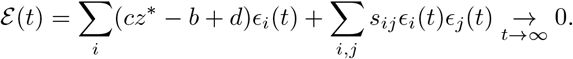

For any *ε* > 0, as soon as *t* is large enough, we deduce that

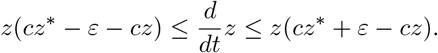

Using classical results on logistic equations, we deduce that *z*(*t*) converges to *z*^∗^. This ends the proof of Proposition 2.1.

### B.3 Complete study of the three allele case

Consider three alleles *A*_1_, *A*_2_, and *A*_3_ whose selection matrix is given by

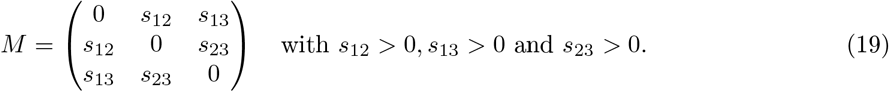

Since the selective advantage parameters are positive, det(*M*) > 0 and one obtains with a simple computation that the condition for existence of a three-allele positive equilibrium *M* ^−1^**1** > 0 can be written as (8). Moreover, the three-alleles equilibrium equals

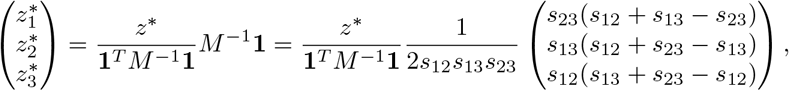

where

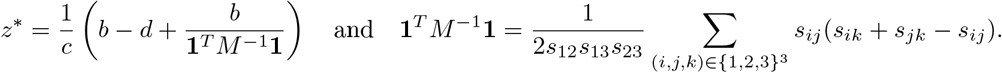

In this case, the behavior of the dynamical system (2) can be fully characterized.

#### Proposition B.3

*Let us consider a system of three alleles, whose interactions are characterized by matrix* (19).

- *Assume* (8), *then the co-existence equilibrium given above is globally stable on* 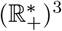.
- *otherwise, assume for example that s*_23_ ≥ *s*_12_ + *s*_13_, *then the two-alleles equilibrium*

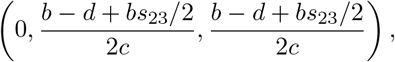

*is globally stable on* 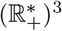.

*Proof.* The first point easily derives from Proposition 2.1.

For the second point, we exhibit a Lyapunov function, which is similar to the one of the previous proof.

Let us consider

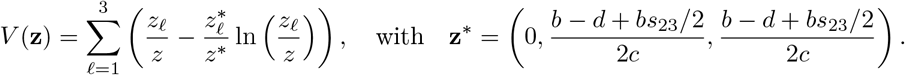

Then, by a direct computation, we find

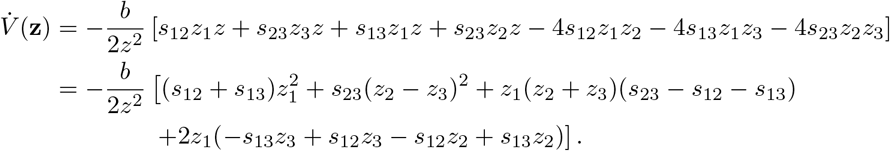

Recall that *s*_23_ ≥ *s*_12_ + *s*_13_, and without loss of generality, we may assume that *s*_12_ ≤ *s*_13_ (otherwise, exchange the roles of *z*_2_ and *z*_3_ in the computations). Then

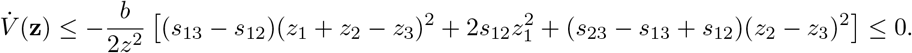

Therefore *V* is a Lyapunov function and 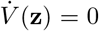 if and only if *z*_1_ = 0 and *z*_2_ = *z*_3_. We then conclude with Lasalle Theorem (Theorem 1 of [14]) and an argument similar to the one at the end of the previous proof.

### B.4 Successive introductions of types

Assume that *k* types coexist. In view of previous sections, if det(*M*) ≠ 0, there is coexistence of the *k* types if and only if *M* ^−1^**1** > 0 and the second eigenvalue of *M* is negative (or equivalently the eigenvalues of *J* ^∗^ are all negative).

If a mutant characterized by parameters *S* = (*s*_*k*+1,*i*_)_*i*=1,..,*k*_ and *σ* = *s*_*k*+1,*k*+1_ appears, this mutant may invade if the Jacobian matrix of (2) around equilibrium (*z*^∗^, 0) is unstable. This Jacobian matrix can be written

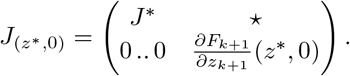

As previously written, the matrix *J* ^∗^ is the jacobian matrix of the resident system of size *k* at equilibrium *z*^∗^ and its eigenvalues are all negative, thus the equilibrium (*z*^∗^, 0) is unstable if the last eigenvalue is positive, *i.e.*

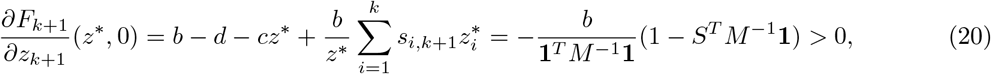

which gives (9).

In what follows, we will be interested in proving the following lemma which states that if the mutant may invade, the new community will be composed of *k* + 1 types if and only if the coexistence state with *k* + 1 types exists.

#### Proposition B.4

*Let us consider a stable resident population with k types given by a matrix M, i.e M satisfies M* ^−1^**1** > 0 *and its second eigenvalue is negative*.

*Let us consider a mutant type arising in this resident population characterized by S and σ. Denote the new fitness matrix by*

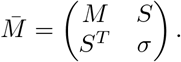

If 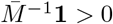, *i.e. the equilibrium with k* + 1 *species exists, then it is globally asymptotically stable*.

*Proof.* Let us first give conditions under which the second eigenvalue of 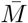 is positive. As 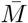 is a symmetric matrix and *M* a principal sub-matrix of 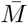, Proposition D.1 implies that

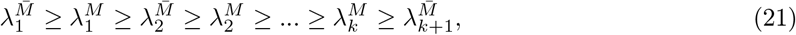

where 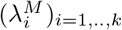 denote the eigenvalues of *M* and 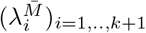 the ones of 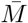. Since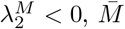 can have at most 2 positive eigenvalues. On the other hand, using Proposition D.2, we compute

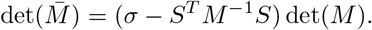

Since det(*M*) has the sign of (−1)^*k*−1^, we deduce the following equivalences:

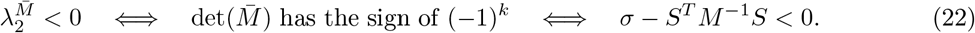

Let us now prove that this last condition holds as soon as 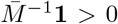. For all *i* ∈ {1, .., *k* + 1} we denote by *M_i_* the matrix resulting from the suppresion of the i*th* column and the i*th* line of 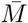, by *S_i_* the i*th* column of 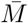 without the i*th* coefficient, and by *σ_i_* the coefficient 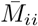. Notice that *M*_*k*+1_ = *M*, *s*_*k*+1_ = *S* and *σ*_*k*+1_ = *σ*. Using these notations and Proposition D.2, we notice that for all *i*

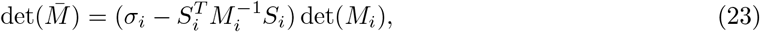

Let us now denote 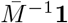 by (*u_i_*)_*i*=1,..,*k*+1_. Then using Cramer’s rule along with (23), we have for all *i* ∈ {1, .., *k* + 1}

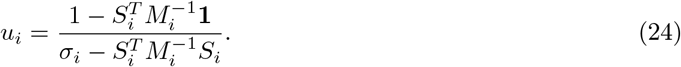

Hence, we deduce that if 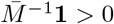, then *u*_*k*+1_ > 0, and since 1 − *S^T^ M* ^−1^**1** < 0, then *σ* − *S^T^ M* ^−1^*S* < 0. Therefore according to (22) and Proposition 2.1, the equilibrium with *k* + 1 species is globally stable. This ends the proof.

Now, let us study the case where the equilibrium with *k* + 1 species does not exist, i.e. there exists *i* ∈ {1, .., *k* + 1} such that *u_i_* < 0. We are not capable of deciphering which alleles will go extinct due to mutant invasion. Numerical simulations reveal that alleles 1 ≤ *i* ≤ *k* that verify *u*_*i*_*u*_*k*+1_ > 0 might go extinct in the equilibrium. Notice that in some cases, all these alleles disappear, while in other cases, only a subset does.

## C Proofs concerning the genetic model

This section gathers theoretical results associated with the specific model in Section 3.1.

### C.1 Globally stable equilibrium in a particular case

In this section we focus on the case where *L* = 3, such that there are 8 possible alleles (see Figure 6 for a representation).

**Figure 6:**
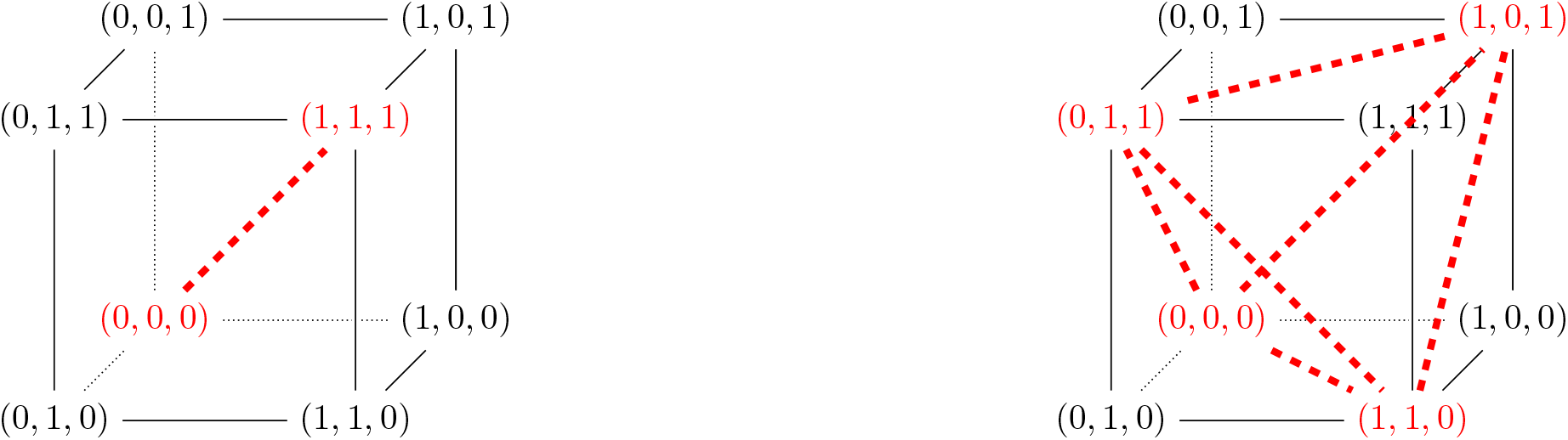
Distances between alleles when assuming 3 sites within the locus *A*, with all possible distances on the edges of the cube, and edges in red showing the alleles maintained at stable state of the population. An example of a population with two most-differentiated alleles is shown on the left (stable population 2), while a example of a stable population with 4 equidistant alleles is represented on the right (stable population 2).

**Figure 7:**
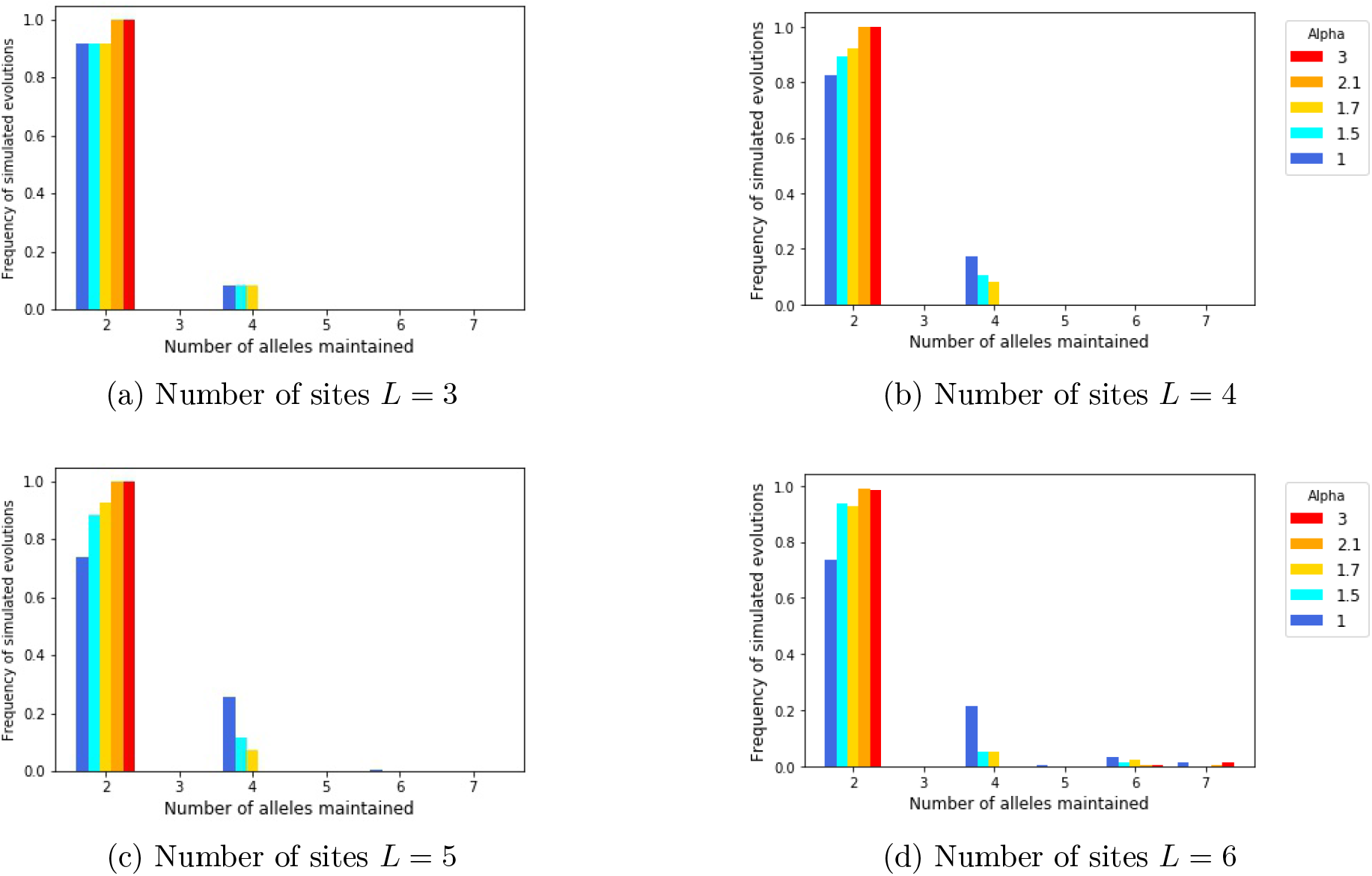
Number of alleles maintained after successive mutations, for different numbers of sites *L* within the locus *A*, and different shapes *α* of the *Genotype-to-reproductive advantage* function. Each graph corresponds to a different number of sites *L* and the color gradient from blue to red indicates increasing values of *α*.

We first prove the convergence of the population state in a particular case for which there is no positive stable equilibrium. This example will be important for the proof of Proposition 3.1 below.

Assume that 4 alleles are present in the population:

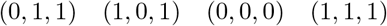

up to permutation of the sites. Then, the selection matrix *M* equals

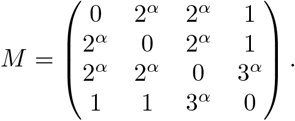

Our aim is to prove that the equilibrium

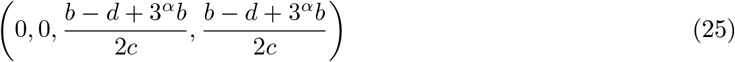

is globally positively stable in (ℝ^+^)^4^, i.e. all trajectories starting from (ℝ^+^)^4^ converge to this equilibrium. To this aim, let us compute the function *V* defined in Equation (14) with *z*^∗^ defined by (25), i.e. 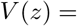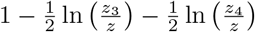. We get

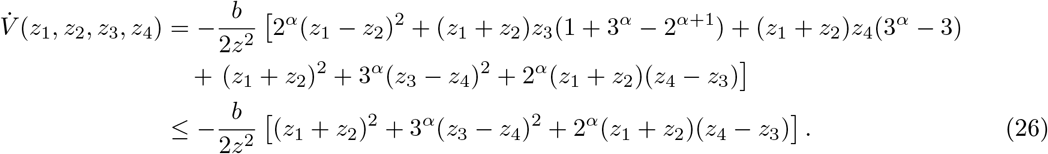

Finally, note that if *α* ≤ 2 then 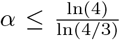, and then the two dimensional function (*x, y*) ↦ *x*^2^ + 3^*α*^*y*^2^+2^*α*^*xy* is positive on ℝ^2^. So *V* is a Lyapunov function and its derivative is null only if *z*_1_ = *z*_2_ = 0 and *z*_3_ = *z*_4_. We then conclude by using an argument similar to the one of the end of proof of Proposition B.2. This finally proves the convergence of all positive trajectories to Equilibrium (25).

### C.2 Proof of Proposition 3.1

Proposition 3.1 arises from two facts: we are able to exhibit stable set of alleles (see Figure 6) and we can characterize the impact of mutations on the number of coexisting alleles in the population.

In fact, we obtain a stochastic process which jumps across stable states of the population depending on the arrival of mutant alleles. This process is characterized by a transition matrix which explicits the possible transitions and is represented on Figure 8.

**Figure 8:**
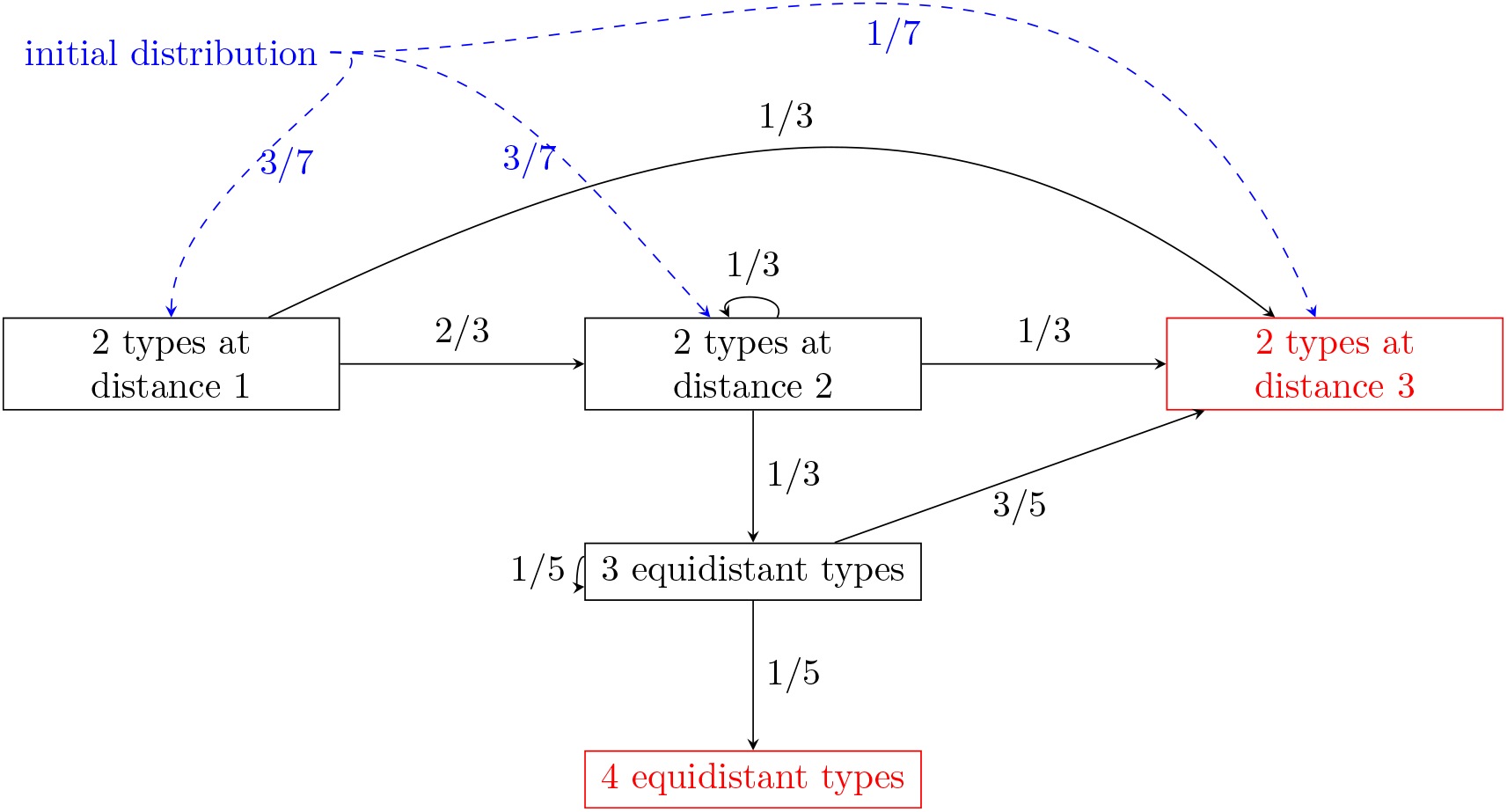
Transition matrix for the allelic composition of the population. The red color corresponds to stable population states. Blue dashed arrows show the initial distribution after the first mutation.

*Proof.* The set of possible alleles is {0, 1}^3^. We first prove that the two following types of genetic compositions are stable: (Stable population 1) Only the two most-differentiated alleles, (Stable population 2) four alleles that are all at distance 2 from each other.

We already know that (stable population 1) is stable from the end of Section 3.1. For (stable population 2), we study the conditions for invasion of a mutant in a population composed of four alleles all at distance 2 from each other. The new mutant will necessary be at distance 1 from 3 of the resident types, and at distance 3 from the forth one. Then following the notations of Equation (9) in Section 2.3, we get that *S* = (1, 1, 1, 3^*α*^)^*T*^ . Therefore the new mutant can invade only if

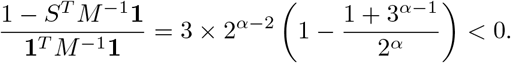

In particular for *α* ∈ [1, 2], this gives that the mutant cannot invade, hence the population with four equidistant alleles is stable.

Now, we consider the different possible orders of mutations arising in the population to prove the proposition. Let us assume without loss of generality, that the population starts with only individuals with allele (0, 0, 0). The second arising allele in the population can be at distance 1 (with probability 3/7), 2 (with probability (3/7), or 3 (with probability 1/7), from (0, 0, 0). In any case, Proposition B.3 implies that this mutant will invade and the population will then exhibit two different alleles at equilibrium.

- If this first mutant is at distance 3 from the resident allele (0, 0, 0), then the population has reached the configuration (Stable population 1) and no new mutation will be able to invade.
- If the first mutant is at distance 2 from the resident allele (0, 0, 0), then there are 3 possibilities for the second mutant (each arising with probability 1/3):

i. The second mutant can be at distance 1 from each of the two pre-existing alleles. Then Equation (9) shows that it cannot invade, if *α* > 1. The population thus stays with the two resident alleles, at a distance 2 from each other.
ii. it can be at distances 1 and 3 from the two first alleles, in which case only the mutant and its opposite type will remain in the population (according to Proposition B.3 and since 3^*α*^ > 2^*α*^ + 1), which corresponds to a configuration (Stable population 1).
iii. The second mutant can be at distance 2 from the two pre-existing alleles. From Equation (8), the population will then converge to an equilibrium with the three first alleles. If the third mutant is in the fourth position at distance 2 from the three pre-existing alleles then it will also invade (see Section 2.3). This mutant arises with probability 1/5 and brings the population to (Stable population 2) (as illustrated in Fig. 6). If the third mutant is in a position such that two most-differentiated alleles co-exist in the population then only these two alleles will remain (see Section C.1 for details). This occurs with probability 3/5. Lastly, if (with probability 1/5) the third mutant is at distance 1 from the three first types then it cannot invade, according to Equation (9).
- Finally, if the first mutant is at distance 1 from (0, 0, 0), then there are two possibilities for the second mutant:

i. The second mutant is at distance 2 and 3 from the two resident alleles, respectively (with probability 1/3). Only the two most-differentiated alleles will then remain from Proposition B.3 and this population state is stable.
ii. The second mutant is at distances 1 and 2 (with probability 2/3) from the pre-existing alleles, respectively. Thus, one of the two pre-existing alleles will disappear and the population will exhibit only two alleles, at distance 2 from each other, as already proved by Proposition B.3.

Finally, the transition graph of the genetic composition of the population is given in Figure 8. This gives that the probability that the population ends up with 4 equidistant genotypes is equal to

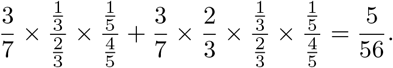

## D Complements of linear algebra

### Proposition D.1 (Eigenvalue Interlacing Theorem)

*Let A* ∈ ℝ^*n*×*n*^ *be a symmetric matrix and B be a principal sub-matrix of size m < n. If* 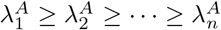 *the eigenvalues of A and* 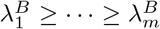 *the eigenvalues of B. Then for any* 1 ≤ *k* ≤ *m*

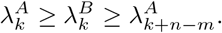

*In particular if m* = *n* − 1 *then*

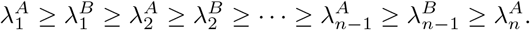

### Proposition D.2 (Schur complement)

*Let M a matrix defined by blocs as* 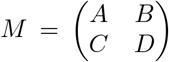 *If D is invertible then the complement of schur of D is defined by A* − *BD*^−1^*C. Moreover*

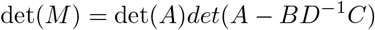

For proof of these two results, see Example 7.5.3 and Section 6.2 in [21].

## Notes

https://plmlab.math.cnrs.fr/costa2150/heterozygote_advantage

### Competing Interest Statement

The authors have declared no competing interest.

